# *Dyrk1a* gene dosage controls bipolar cell development and retinal connectivity

**DOI:** 10.64898/2026.03.15.710015

**Authors:** Christoffer Nord, Amol Tandon, Anthi-Styliani Makiou, Henri Leinonen, Sara Ivy Wilson, Leif Carlsson, Iwan Jones

## Abstract

Mammalian retina development is governed by a core set of eye-field transcription factors that orchestrate tissue lamination and neuronal connectivity through tightly regulated genetic hierarchies. The LIM homeodomain gene *Lhx2* functions as a master eye-field regulator and this study identifies *Dyrk1a* as a previously unrecognised effector of *Lhx2*-dependent programs during mouse retinal development. Conditional deletion of *Dyrk1a* in retinal progenitor cells led to increased apoptosis within the inner retina demonstrating a requirement for *Dyrk1a* in cell survival and lineage maintenance. Adult heterozygous mice exhibited dorsoventral reductions in bipolar cell number and indicated a gene-dosage-sensitive control of interneuron survival. Bipolar cell loss disrupted their mosaic organisation and resulted in diminished activity across both scotopic and photopic retinal pathways. *Dyrk1a* haploinsufficient mice also displayed pronounced disruption of inner plexiform layer stratification consistent with defective laminar targeting and synaptic partner integration. This study defines a gene-dosage-dependent mechanism through which transcriptional hierarchies regulate neuronal connectivity and function during mammalian retinal development.

## Introduction

Development of the mammalian retina involves the regulated progression of retinal progenitor cells (RPCs) through a series of temporospatial differentiation waves to generate six principle neuronal classes: rod and cone photoreceptors, horizontal, bipolar, amacrine and retinal ganglion cells (RGCs) and Müller glia. In the adult retina these lineages are arranged into a laminated structure comprised of three nuclear layers and two intervening plexiform layers. Photoreceptors occupy the (i) outer nuclear layer (ONL), (ii) bipolar, horizontal and amacrine cells in addition to Müller glia reside in inner nuclear layer (INL) while (iii) RGCs and displaced amacrine cells are localised to the ganglion cell layer (GCL). Synaptic connectivity between photoreceptors, bipolar and horizontal cells occur in the outer plexiform layer (OPL) whereas bipolar, amacrine and RGCs form defined stratification patterns within the inner plexiform layer (IPL). Beyond this radial organisation the neuronal classes within each nuclear layer (ONL, INL and GCL) are also precisely arranged into well-ordered tangential mosaics that ensure uniform sampling of visual space (Amini et al., 2017; Reese and Keeley, 2015).

Retinal development is governed by a core group of eye-field transcription factors (EFTFs) (Diacou et al., 2022) that orchestrate both temporospatial RPC differentiation and subsequent lamination and synaptic connectivity of the principle neuronal classes (Esteve and Bovolenta, 2006; Shiau et al., 2021). Central to this process is the LIM homeobox transcription factor *Lhx2* (de Melo et al., 2018; Gordon et al., 2013; Hagglund et al., 2011) which exerts transcriptional control over downstream genetic hierarchies by forming protein complexes with cofactors such as *Ldb1* and *Ldb2* (Gueta et al., 2016). Given the complexity of retinal development many direct transcriptional targets or genes involved in *Lhx2*-driven regulatory hierarchies remain unidentified. Of particular interest are genes governing apoptosis and tangential mosaic formation as these processes are essential for correct retinal organisation and tissue-wide connectivity (Dhomen et al., 2006). This knowledge gap also raises a central question in developmental biology: how do gate-keeper transcriptional regulators orchestrate their effects to generate diverse cell types and neural connectivity?

To address this open question the present study used a bioinformatics approach to uncover novel transcriptional targets of *Lhx2* during retinal development. Analysis of publicly available RNA-sequencing datasets (de Melo et al., 2016; Li et al., 2022) revealed *Dyrk1a* as a previously unrecognised downstream target gene. *Dyrk1a* is a member of the evolutionary conserved dual specificity tyrosine-phosphorylation-regulated kinase family (*DYRK*) (Arranz et al., 2019; Hammerle et al., 2011). *Dyrk1a* gene-dosage is an important regulator of nervous system development (Arbones et al., 2019; Hammerle et al., 2011) since haploinsufficiency leads to aberrant corticogenesis whereas overexpression impairs axonal growth and synaptogenesis (Martinez de Lagran et al., 2012). Moreover, trisomy of the human *DYRK1A* gene is linked to Down syndrome (OMIM #190685) while mutations are associated with intellectual developmental disorder (MRD7) (OMIM #614104) (Johnson et al., 2025).

Despite ocular abnormalities being reported in MRD7 individuals (Cai et al., 2023; Mejecase et al., 2021) the specific roles of *Dyrk1a* during retina development remain largely unexplored. Mouse models have demonstrated that germline *Dyrk1a* haploinsufficiency leads to a reduction in cell number while *Dyrk1a* gain-of-function leads to increased nuclear layer thickness (Laguna et al., 2008; Laguna et al., 2013). While these studies demonstrate that *Dyrk1a* gene-dosage is essential for retinal development the use of germline models cannot resolve whether the observed phenotypes arise from *Dyrk1a* cell-autonomous functions. To address this question this study describes the generation of conditional mouse models that delete *Dyrk1a* in RPCs during retinogenesis. Loss of one of both *Dyrk1a* alleles resulted in increased programmed cell death and neonatal lethality in homozygous animals and inner retina abnormalities in heterozygous mice. This phenotype in *Dyrk1a* haploinsufficient mice was best characterised by a dorsoventral reduction in interneuron number with bipolar cells being particularly affected. Region-specific deviations in bipolar cell densities disrupted their mosaic arrangement and resulted in reduced bipolar cell-driven retinal activity. Heterozygous mice also displayed pronounced disruption of inner plexiform layer stratification consistent with defective laminar targeting and synaptic partner integration. This study therefore identifies *Dyrk1a* as a critical effector of *Lhx2*-regulated transcriptional hierarchies and provides new insights into the gene-dosage-dependant roles of *Dyrk1a* during bipolar cell mosaic formation and inner retina connectivity.

## Results

### Identification of Dyrk1a as a target gene of Lhx2*-*mediated transcriptional programs

To identify target genes of *Lhx2*-regulated transcriptional hierarchies a bioinformatics workflow was developed to interrogate publicly available RNA-sequencing datasets (GSE172457 and GSE75889) (de Melo et al., 2016; Li et al., 2022). These datasets were derived from RPCs isolated from control and *Lhx2* conditional knockout (cKO) mice during embryonic (E) and postnatal (P) development (E15.5 and P2) (Figure 1A). The rationale behind this workflow was that genes consistently downregulated at both developmental ages were likely core components of *Lhx2*-regulated programs operating throughout critical phases of retinal development. DESeq2 analysis (padj <0.05, baseMean >10, log2FC <-0.3) identified 1549 genes that were significantly downregulated in cKO retinas at E15.5 and 147 transcripts with reduced expression at P2 (Figure 1A). These differentially expressed genes (DEGs) were intersected to identify 117 core genes that exhibited decreased expression at both developmental ages (Figure 1A – 1B). Gene ontology (GO) analysis of this core dataset revealed significant enrichment for genes known to be involved in visual system development (e.g. *Vsx2* and *Six6*) and cell cycle regulation (e.g. *Cdkn3* and *Ccnd1*) (Figure 1C). These findings supported the idea that *Lhx2*-regulated transcriptional hierarchies orchestrate retinal development (de Melo et al., 2018; Gordon et al., 2013; Hagglund et al., 2011) and validated that our bioinformatics workflow effectively identified *Lhx2*-regulated target genes.

**Figure 1.**
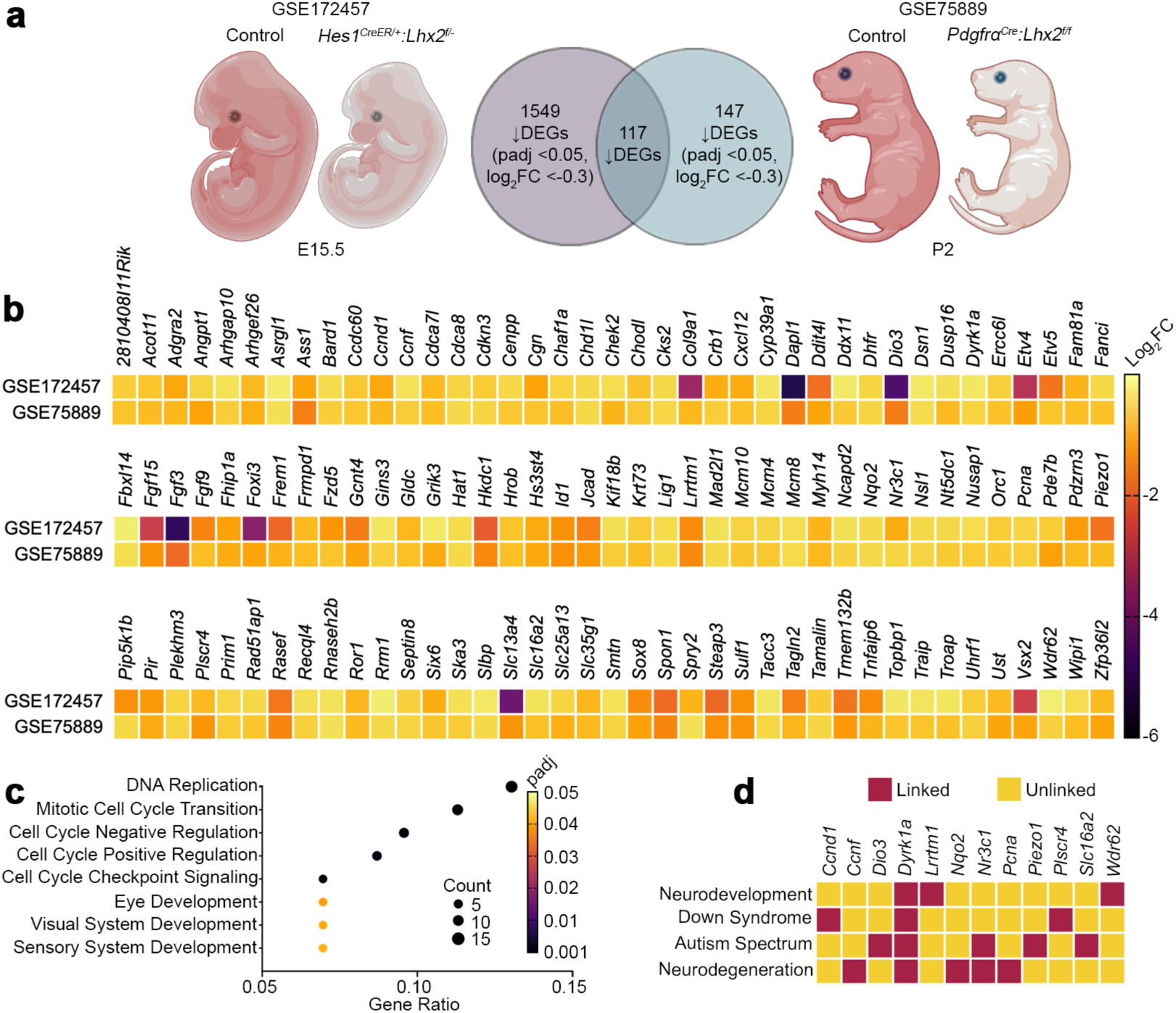
Identification of *Dyrk1a* as a target of *Lhx2*-mediated transcriptional programs. (A) Differential expression analysis of RNA-sequencing datasets (GSE172457 and GSE75889) derived from the retinas of control and *Lhx2* cKO mice. A total of 1549 genes were significantly downregulated in cKO animals at E15.5 while 147 transcripts showed reduced expression at P2. (B) Heatmap analysis of intersected DEGs demonstrate that these genes exhibit decreased expression in cKO animals at both developmental ages. (C) Gene ontology (GO) analysis of the intersected genes demonstrates a significant enrichment for biological processes related to visual system development and cell cycle regulation. (D) Several intersected DEGs are linked to a broad range of human nervous system disorders. Created in BioRender. Biorender.com/xxmk16m.

### Temporospatial expression of Dyrk1a during mouse eye development

Through bioinformatic analysis *Dyrk1a* was identified as an *Lhx2*-regulated target gene (Figure 1B) and given its involvement in human CNS disorders (Figure 1D) represented a compelling candidate for further study. Characterisation of *Dyrk1a* temporospatial distribution during mouse retinal development (Figure S1) demonstrated that transcripts were broadly distributed in the retina throughout embryonic development (E10.5 – E18.5, Figure S1A – S1E) and became enriched in the outer neuroblastic layer (ONBL), inner neuroblastic layer (INBL) and GCL at later ages (E16.5 - E18.5, Figure S1D – S1E, arrowheads). Protein distribution generally mirrored embryonic gene expression (Figure S1F – S1J). Dyrk1a was initially observed in the ventral retina at E10.5 (Figure S1F, arrowhead) but subsequently became enriched throughout the retina from E12.5 to E14.5 (Figure S1G – S1H, arrowheads). Dyrk1a then increased throughout the ONBL, INBL and GCL at later ages (E16.5 – E18.5, Figure S1I – S1J, arrowheads). During postnatal development *Dyrk1a* expression and protein became progressively restricted to specific retinal layers (Figure S1K – S1T). *Dyrk1a* transcripts and immunoreactivity were initially observed within the INBL and GCL at birth (Figure S1K and S1P, arrowheads) and accumulated in the apical INL and GCL at P7 (Figure S1L and S1Q, arrowheads). At later postnatal ages (P14, P21 and P42) *Dyrk1a* mRNA and protein became restricted to the photoreceptor segments and inner retina (INL and GCL) (Figure S1M – S1O and S1R – S1T, arrowheads). Collectively, the dynamic temporospatial pattern of *Dyrk1a* activity suggested its involvement in molecular programs that regulated embryonic RPC differentiation as well as postnatal lamination and mosaic organisation of the inner retina.

### Conditional deletion of Dyrk1a leads to genotype-dependent allelic recombination

Conditional mouse models were generated to investigate the roles of *Dyrk1a* during retinal development. Animals carrying floxed *Dyrk1a* alleles (*Dyrk1a^tm1Jdc/J^*, referred to as *Dyrk1a^+/f^* or *Dyrk1a^f/f^*) (Thompson et al., 2015) were crossed with mice harbouring an *Lhx2* promoter-driven *Cre* recombinase transgene (*Tg(Lhx2-Cre)1Lcar*, referred to as *Lhx2-Cre*). This transgene induced recombination of the *Dyrk1a* allele in RPCs during optic vesicle-to-optic cup transition (E8.5–E12.5) (Hagglund et al., 2011) and its dorsoventral activity profile (Jones et al., 2019) generated a spatial gradient of reduced *Dyrk1a* dosage across the retina. Mutant mice were born at normal Mendelian ratios (Table S1) but homozygous animals died shortly after birth due to recombination of both *Dyrk1a^tm1Jdc/^*^J^ alleles within CNS regions essential for neonatal survival (Hagglund et al., 2011).

Given the splicing complexity of mouse *Dyrk1a* gene (*ENSMUSG00000022897*) (Figure 2A) a combination of quantitative PCR (qPCR) and immunoblotting was employed to evaluate allele recombination efficiency and protein levels, respectively. Tissue was harvested at P0 for these analyses since robust *ROSA26R^R/R^*reporter activity indicated widespread *Cre*-mediated recombination in all genotypes (Figure 2B – 2E). Significant genotype-dependent reductions in *Dyrk1a^tm1Jdc/^*^J^ allele (Figure 2F) and protein levels (Figure 2G – 2H and Figure S2) were observed in heterozygous and homozygous animals relative to transgenic controls. In addition, cleaved Caspase3 levels were significantly increased in *Lhx2-Cre:Dyrk1a^f/f^*retinas (Figure 2G, 2I and Figure S2) consistent with the established role of *Dyrk1a* as a regulator of programmed cell death during retinal development (Laguna et al., 2008). Together, this data confirmed efficient conditional deletion of *Dyrk1a* and established the suitability of this mouse model for investigating its role during retinal development.

**Figure 2.**
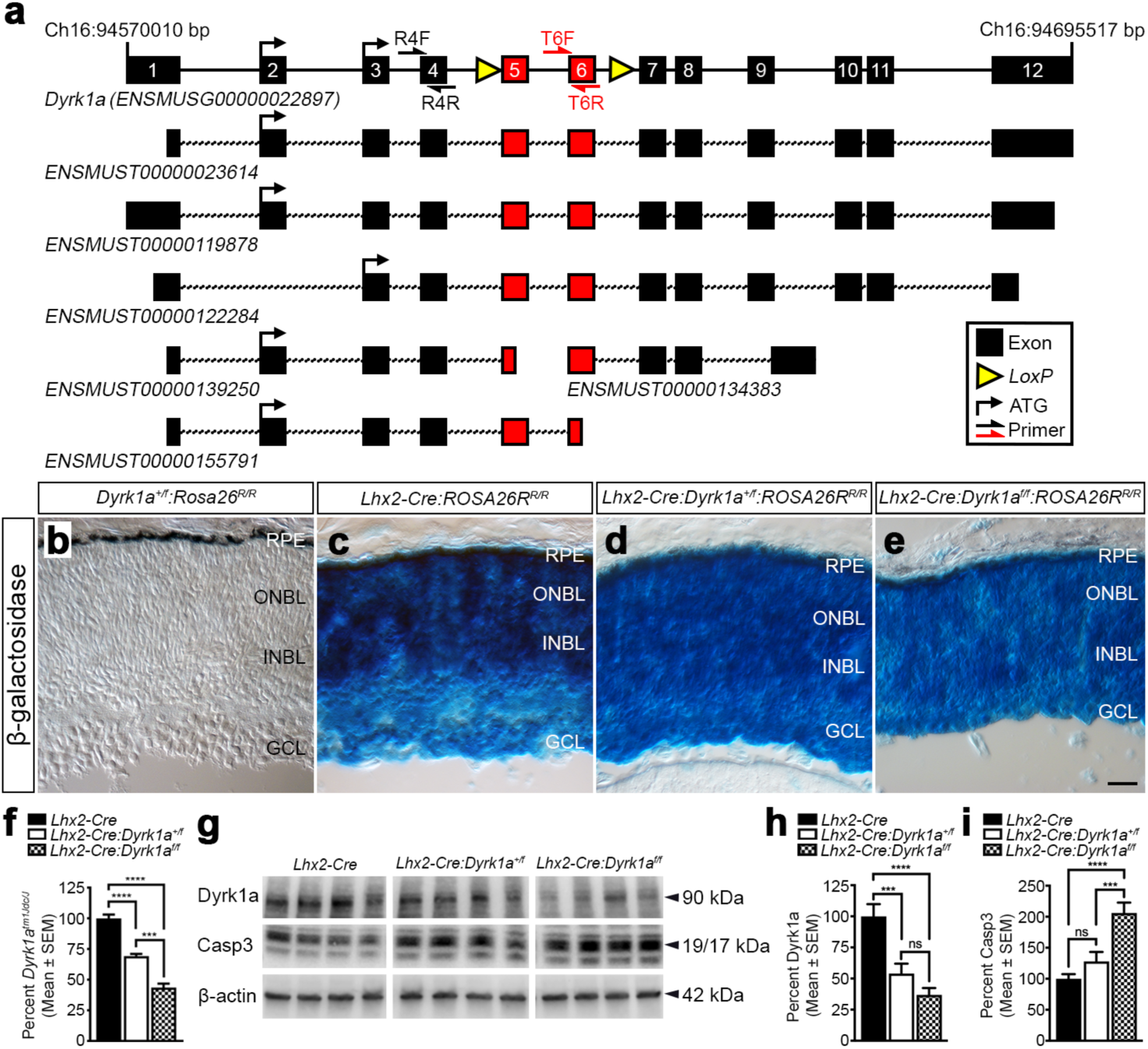
Conditional deletion of *Dyrk1a* leads to genotype dependent allelic recombination. (A) Schematic diagram illustrating the genomic organisation of the *Dyrk1a^tm1Jdc/J^*allele (*ENSMUSG00000022897*, Chr16:94570010 – 94695517 bp). The gene comprises 12 coding exons (black boxes) and gives rise to six mRNA isoforms. The longest transcripts (*ENSMUST00000023614*, *ENSMUST00000119878, ENSMUST00000122284)* encode a protein of approximately 90 kDa. *LoxP* sites (yellow triangles) flank exons 5 and 6 and facilitate *Cre*-mediated recombination of the kinase domain in the translated protein. Oligonucleotides used to assess the extent of allele recombination were designed to amplify across reference exon 4 (R4F and R4R, black arrows) and target exon 6 (T6F and T6R, red arrows). (B – E) Representative lineage tracing analysis of *Dyrk1a^+/f^:ROSA26R^R/R^*(B), *Lhx2-Cre:ROSA26R^R/R^* (C), *Lhx2-Cre:Dyrk1a^+/f^:ROSA26R^R/R^*(D) and *Lhx2-Cre:Dyrk1a^f/f^:ROSA26R^R/R^* (E) at P0. Robust β-galactosidase activity is detected throughout the retina of all animals that carry the *Lhx2-Cre* transgene (C – E) confirming *Cre*-mediated recombination of the *ROSA26R^R/R^* reporter allele. (F) Quantitative analysis of *Dyrk1a^tm1Jdc/J^* allele recombination efficiency in genomic DNA harvested from the retina of *Lhx2-Cre* (*n* = 4), *Lhx2-Cre:Dyrk1a^+/f^* (*n* = 4) and *Lhx2-Cre:Dyrk1a^f/f^*(*n* = 4) mice at P0. A significant genotype-dependent loss of the *Dyrk1a LoxP* region was observed in heterozygous and homozygous animals, respectively. Quantification is based on the level of product generated by the *LoxP* target primer pair normalised against the level of product amplified by the reference primer pair for each biological replicate. (G) Representative immunoblots showing the main 90 kDa Dyrk1a band and the 19/17 kDa cleaved Caspase3 bands with β-actin as a normalisation control in soluble protein extracts prepared from the retina and pigment epithelium of *Lhx2-Cre*, *Lhx2-Cre:Dyrk1a^+/f^*and *Lhx2-Cre:Dyrk1a^f/f^* mice at P0. (H) Quantitative analysis of Dyrk1a levels in soluble protein extracts prepared from the retina and pigment epithelium of *Lhx2-Cre* (*n* = 12), *Lhx2-Cre:Dyrk1a^+/f^*(*n* = 12) and *Lhx2-Cre:Dyrk1a^f/f^* (*n* = 12) mice at P0. Dyrk1a levels were significantly reduced in a genotype-dependent manner in heterozygous and homozygous animals. Quantification is based on the 90 kDa Dyrk1a protein band normalised to β-actin within the same biological replicate. (I) Quantitative analysis of cleaved Caspase3 levels in soluble protein extracts prepared from the retina and pigment epithelium of *Lhx2-Cre* (*n* = 12), *Lhx2-Cre:Dyrk1a^+/f^* (*n* = 12) and *Lhx2-Cre:Dyrk1a^f/f^*(*n* = 12) mice at P0. A genotype-dependent increase in cleaved Caspase3 levels was observed in heterozygous mice with significance reached in homozygous animals. Quantification is based on the 19/17 kDa cleaved Caspase3 protein band normalised to β-actin in the same biological replicate. All data represents the mean ± SEM values. Statistical differences were calculated using one-way ANOVA followed by a Tukey’s multiple comparison test. p-values are denoted as follows: ***p ≤ 0.001 and ****p ≤ 0.0001. Scale bar: (B – E) 50 μm. Abbreviations: bp, base pair; Casp3, cleaved Caspase3; Ch16, chromosome 16; GCL, ganglion cell layer; INBL, inner neuroblastic layer; ONBL, outer neuroblastic layer; R, reference primer pair; RPE, retinal pigment epithelium; T, target primer pair.

### Conditional deletion of Dyrk1a impairs retinal neurogenesis through apoptotic deficits

Given that increased apoptosis was observed in mutant animals the effect of *Dyrk1a* loss on RPC survival was investigated. The numbers of amacrine and RGCs were quantified as representative neuronal subtypes broadly distributed in the retina during embryonic (E14.5) and postnatal development (P0) (Figure 3). A genotype-dependent reduction in the number of Isl1/2^+^ cells (labelling both amacrine and RGCs) (Elshatory et al., 2007) was observed in the retina of *Lhx2-Cre:Dyrk1^+/f^* and *Lhx2-Cre:Dyrk1a^f/f^*mice (Figure 3A – 3F) although statistical significance was only reached in homozygous animals (Figure 3G – 3H). Cleaved Caspase3 immunostaining was subsequently performed to assess the contribution of programmed cell death to this reduction in neuron number. A genotype-dependent increase in the number of apoptotic cells was detected within the retina of both mutant models (Figure 3A – 3F, arrowheads in inset images) with statistical significance again observed in homozygous animals (Figure 3I – 3J). These findings indicated that *Dyrk1a*-ablation during retinal development compromised neuronal survival through apoptotic mechanisms in a gene dosage-dependant manner.

**Figure 3.**
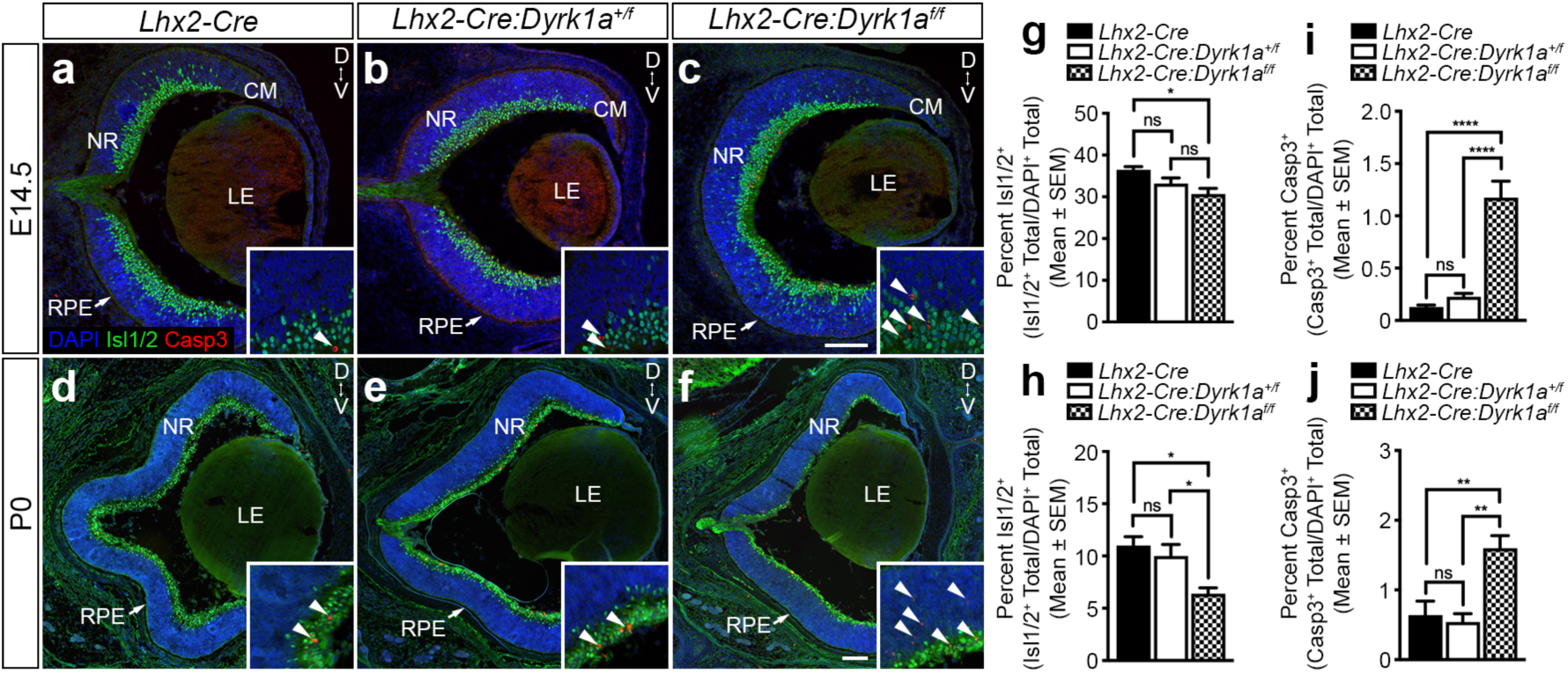
Condition deletion of *Dyrk1a* impairs retinal neurogenesis through apoptotic deficits. (A – C) Representative coronal sections taken from the retina of *Lhx2-Cre* (A), *Lhx2-Cre:Dyrk1a^+/f^* (B) and *Lhx2-Cre:Dyrk1a^f/f^* (C) at E14.5 to illustrate the presence of Isl1/2^+^ neurons and cleaved Caspase3^+^ apoptotic cells (A – C, arrowheads in inset images). (D – F) Representative coronal sections taken from the retina of *Lhx2-Cre* (D), *Lhx2-Cre:Dyrk1a^+/f^* (E) and *Lhx2-Cre:Dyrk1a^f/f^* (D) at P0 to illustrate the presence of Isl1/2^+^ neurons and cleaved Caspase3^+^ apoptotic cells (D – F, arrowheads in inset images). (G – H) Quantitative analysis of *Lhx2-Cre* (*n* = 5), *Lhx2-Cre:Dyrk1a^+/f^*(*n* = 5) and *Lhx2-Cre:Dyrk1a^f/f^* (*n* = 5) animals at E14.5 (G) and P0 (H) demonstrates a significant reduction in the number of amacrine and RGCs in the retinas of homozygous animals at both ages. (I – J) Quantitative analysis of *Lhx2-Cre* (*n* = 5), *Lhx2-Cre:Dyrk1a^+/f^*(*n* = 5) and *Lhx2-Cre:Dyrk1a^f/f^* (*n* = 5) mice at E14.5 (I) and P0 (J) demonstrates a significant increase the number of apoptotic cells in the retinas of *Lhx2-Cre:Dyrk1a^f/f^* animals at both ages. All data represents the mean ± SEM values. Statistical differences were calculated using one-way ANOVA followed by a Tukey’s multiple comparison test. p-values are denoted as follows: *p ≤ 0.05, **p ≤ 0.01 and ****p ≤ 0.0001. Scale bars: (A – F) 100 μm. Abbreviations: Casp3, cleaved Caspase3; CM, ciliary margin; D, dorsal; LE, lens; NR, neural retina; RPE, retinal pigment epithelium; V, ventral.

### Dyrk1a haploinsufficiency leads to a reduction in inner retina lamination thickness

Due to the neonatal lethality observed in homozygous animals (Table S1) the long-term impact of complete *Dyrk1a* loss on postnatal retinal development could not be determined. Subsequent studies were therefore focussed on *Lhx2-Cre:Dyrk1a^+/f^* mice. These haploinsufficient animals survived to adulthood and provided a model of human MRD7 syndrome where affected individuals exhibit ocular abnormalities (Ji et al., 2015). Moreover, the genotype-dependent increases in apoptosis observed in neonatal *Lhx2-Cre:Dyrk1a^+/f^*retinas (Figure 3G – 3J) suggested that chronic reduction in *Dyrk1a* dosage may be sufficient to disrupt retinal development and visual function in adult mice.

To provide a global overview of laminar organisation the retinas of *Lhx2-Cre* and *Lhx2-Cre:Dyrk1a^+/f^* adult animals (6 – 8 weeks) were assessed by optical coherence tomography (OCT) (Figure 4). The ONL of both groups displayed comparable thickness across all axes (Figure 4A and 4D). However, a significant reduction in INL and IPL thickness was observed in dorsal-associated regions in heterozygous animals (Figure 4B - 4F). To confirm this inner retina phenotype histological analyses were performed across four designated dorsoventral quadrants (D1, D2, V3 and V4) spanning the entire retina relative to the optic nerve head (Figure 4G – 4N). The retina of transgenic control mice displayed a well-ordered laminated structure that was organised into three distinct nuclear layers (ONL, INL and GCL) being intersected by two plexiform layers (OPL and IPL) (Figure 4G – 4J). In contrast, while *Lhx2-Cre:Dyrk1a^+/f^* retinas exhibited comparable lamination a significant decrease in apicobasal thickness was observed across all dorsoventral quadrants (Figure 4K – 4R). Quantification of individual layer thickness demonstrated that this phenotype was primarily driven by a reduction in INL and associated plexiform layer (OPL and IPL) thickness particularly in the dorsal region (Figure 4O – 4R). Taken together, these analyses demonstrated that chronic reduction in *Dyrk1a* dosage induced a strong inner retinal phenotype and indicated that *Dyrk1a* haploinsufficiency disrupted the survival of inner retinal neurons likely due to increased apoptosis.

**Figure 4.**
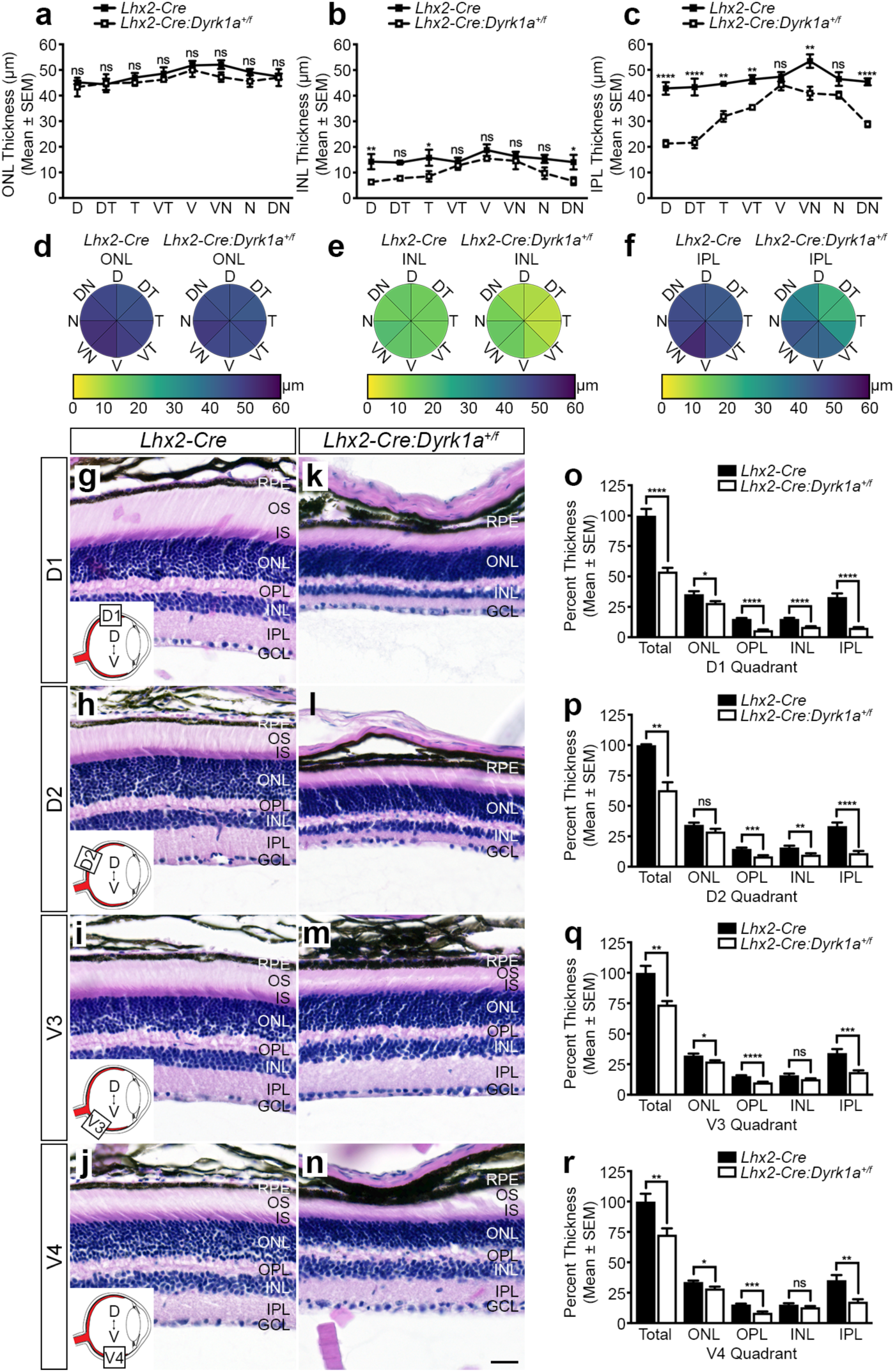
*Dyrk1a* haploinsufficiency leads to reduced inner retina lamination thickness. (A – C) Quantitative optical coherence tomography analysis of ONL (A), INL (B) and IPL (C) thickness taken at cardinal (D, V, N, T) and intercardinal (DN, DT, VN, VT) orientations in the retina of *Lhx2-Cre* (*n* = 3) and *Lhx2-Cre:Dyrk1a^+/f^*(*n* = 3) animals (6 – 8 weeks). All measurements were performed at fixed distance (500 µm) from the optic nerve head. ONL thickness was comparable between both groups (A) while a significant reduction in INL and IPL thickness was predominantly localised to dorsal-associated regions in heterozygous animals (B – C). (D – F) Heat map plots showing layer thickness distribution measured by optical coherence tomography in the ONL (D), INL (E) and IPL (F) of *Lhx2-Cre* and *Lhx2-Cre:Dyrk1a^+/f^* animals. ONL thickness was comparable between both groups (D) while a significant reduction in INL and IPL thickness was spatially localised to dorsal associated regions in heterozygous animals (E – F). (G – N) Representative histological analysis of coronal eye sections taken from *Lhx2-Cre* and *Lhx2-Cre:Dyrk1a^+/f^*adult animals (6 – 8 weeks) across four designated dorsoventral quadrants (D1, D2, V3 and V4) spanning the entire retina relative to the optic nerve head. The retina of *Lhx2-Cre* mice (G – J) displayed a well-ordered laminated structure that was organised into three distinct nuclear layers (ONL, INL and GCL) being intersected by two plexiform layers (OPL and IPL). In contrast, the laminated retina of *Lhx2-Cre:Dyrk1a^+/f^* animals (K – N) exhibited a dorsoventral gradient of reduced apicobasal thickness with the highest penetrance being observed in the dorsal hemisphere (K – L). (O – R) Quantitative assessment of total and individual layer thicknesses across four designated dorsoventral quadrants of *Lhx2-Cre* (*n* = 7) and *Lhx2-Cre:Dyrk1a^+/f^*(*n* = 7) adult animals (6 – 8 weeks). Heterozygous mice exhibited a significant decrease in total retina thickness along with selective reduction of individual layers with the greatest differences being observed in the dorsal INL and associated plexiform layers (OPL and IPL) (O – P). All data represents the mean ± SEM. Statistical differences were calculated using a two-way ANOVA followed by a Sidak’s multiple comparison test (A – C) or unpaired two-tailed Student’s t-test (O – R). p-values are denoted as follows: *p≤ 0.05, **p≤ 0.01, ***p≤ 0.001 and ****p≤ 0.0001. Scale bar: (G – N) 25 μm. Abbreviations: D, dorsal; DN, dorsonasal; DT, dorsotemporal; GCL, ganglion cell layer; INL, inner nuclear layer; IPL, inner plexiform layer; IS, inner segments; N, nasal; ONL, outer nuclear layer; OPL, outer plexiform layer; OS, outer segments; RPE, retinal pigment epithelium; T, temporal; V, ventral; VN, ventronasal; VT, ventrotemporal.

### Dyrk1a haploinsufficiency reduces the number and connectivity of inner retinal neurons

Immunohistochemical analyses using a panel of neuronal markers was performed to establish whether the apoptotic phenotype was truly confined to the inner retina of homozygous animals. The ONL of both transgenic control and heterozygous mice were comparably populated by densely packed photoreceptor cell bodies (DAPI^+^, Figure S3A – S3X) and exhibited well-developed segments enriched in rhodopsin photopigment across all dorsoventral quadrants (Figure S3A – S3H). However, discrete rhodopsin^+^ puncta were observed within the pigment epithelium of transgenic controls consistent with rod outer segment phagocytosis (Figure S3A – S3D, arrows). In contrast, *Lhx2-Cre:Dyrk1a^+/f^*retinas exhibited an absence of rhodopsin phagosomes in the dorsal region (Figure S3E – S3F, asterisks) while ventral quadrants appeared comparable to transgenic controls (Figure S3G – S3H, arrowheads). Both genotypes also exhibited characteristic dorsoventral gradients of short-wave (Figure S3I – S3P) and medium-wave (Figure S3Q – S3X) cone opsin expression. These data indicated that photoreceptor structure and integrity was largely preserved in both groups although heterozygous animals displayed a dorsal-specific defect in rod outer segment phagocytosis suggestive of early photoreceptor degeneration (Kevany and Palczewski, 2010).

Inner retina composition differed significantly between *Lhx2-Cre* and *Lhx2-Cre:Dyrk1a^+/f^* animals (Figure 5A – 5X). The INL of transgenic controls was populated by bipolar (Chx10^+^, Figure 5A – 5D, arrowheads), amacrine (Pax6^+^, Figure 5I – 5L, arrowheads) and horizontal (Calbindin^+^, Figure 5Q – 5T, arrowheads) cells with characteristic densities and apicobasal positioning. RGCs and displaced amacrine cells also displayed similar numbers and distribution in the GCL (Pax6^+^, Figure 5I – 5L, arrowheads). In contrast, although similar spatial positioning of these neuronal subtypes was observed in heterozygous animals (Chx10^+^, Figure 5E – 5H; Pax6^+^, Figure 5M – 5P; Calbindin^+^, Figure 5U – 5X, arrowheads) the total number of neurons within the INL and GCL were reduced across all dorsoventral quadrants with the most pronounced deficits occurring within the dorsal-most region (Figure 5E, 5M and 5U). A panel of neurite-specific markers were subsequently employed to determine whether this reduction in inner retina neuron number influenced IPL stratification (Figure 6). Rod ON bipolar cells in transgenic controls exhibited characteristic sublamina stratification (PKCα^+^, S4/S5, Figure 6A – 6D, arrowheads). Likewise, various amacrine cell subtypes showed consistent and well-defined neurite termination patterns. These included starburst amacrine cell arborising in S2/S4 (chAT^+^, Figure 6I – 6L, arrowheads) and All amacrine cells terminating in S1/S2, S2/S3 and S3/S4 (Calretinin^+^, Figure 6Q – 6T, arrowheads). In contrast, homozygous retinas displayed aberrant neurite stratification patterns. The reduced population of rod ON bipolar cells exhibited irregular neurite organisation within the S4/S5 sublamina (PKCα^+^, Figure 6E – 6H, arrowheads). Similarly, starburst (chAT^+^, Figure 6N – 6P, arrowheads) and All amacrine cells (Calretinin^+^, Figure 6U – 6X, arrowheads) displayed disrupted IPL stratification profiles. Collectively, *Dyrk1a* haploinsufficiency resulted in global loss of inner retina neurons along the dorsoventral axis with the most severe reduction in the dorsal retina leading to the greatest IPL stratification deficits.

**Figure 5.**
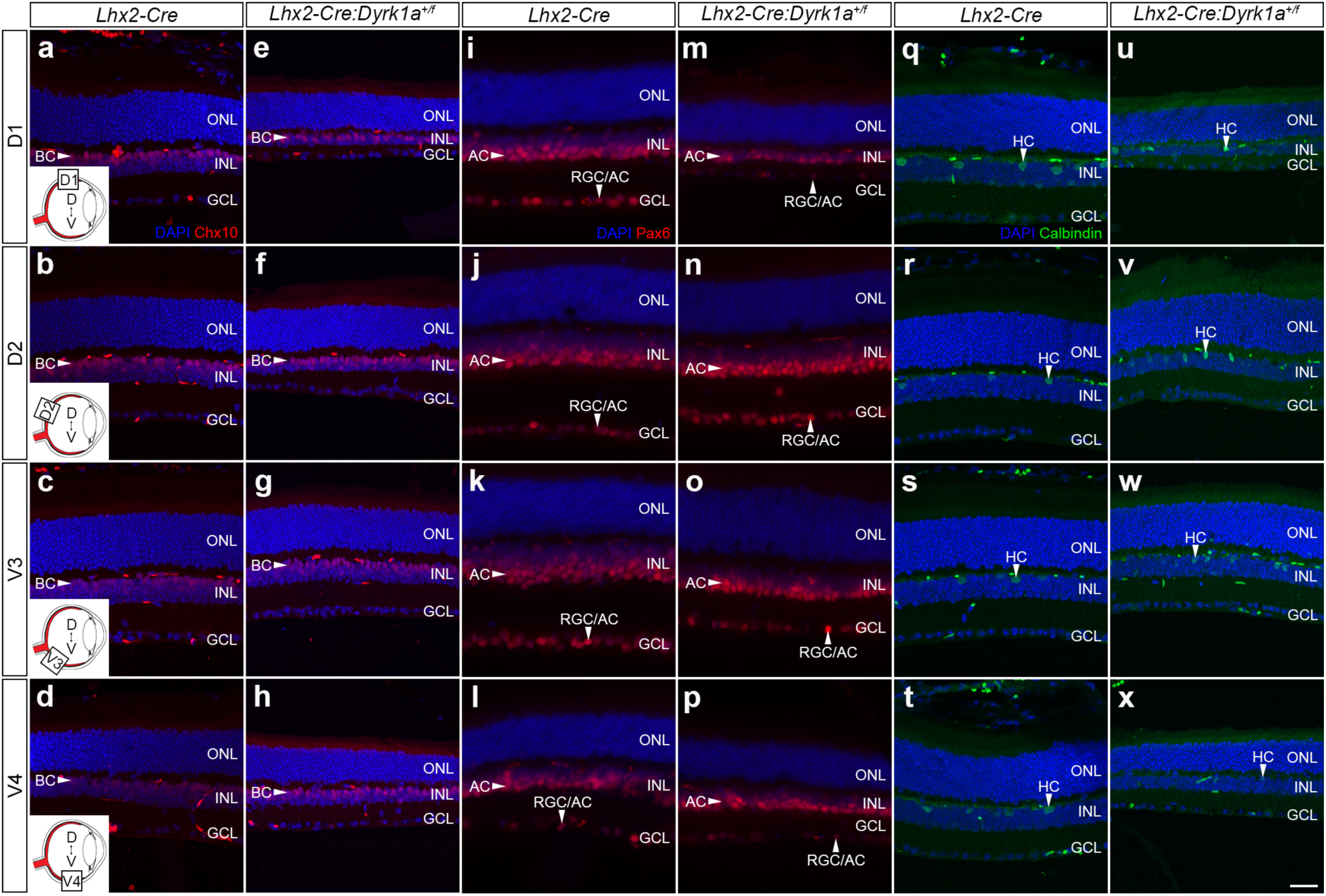
*Dyrk1a* haploinsufficiency reduces inner retina cell number. (A – X) Representative immunohistochemical staining of coronal eye sections taken from *Lhx2-Cre* and *Lhx2-Cre:Dyrk1a^+/f^* adult mice (6 – 8 weeks) across four designated dorsoventral quadrants (D1, D2, V3 and V4) spanning the entire retina relative to the optic nerve head. (A – H) All quadrants in the INL of *Lhx2-Cre* (A – D) animals were comparably populated with bipolar cells (Chx10^+^, BC, arrowheads). In contrast, the dorsoventral gradient of reduced INL thickness in *Lhx2-Cre:Dyrk1a^f/f^* animals (E – H) was accompanied by an apparent reduction in total bipolar cell number (Chx10^+^, BC, arrowheads) which was most prominent in the D1 quadrant (E). (I – P) The INL and GCL of *Lhx2-Cre* animals (I – L) were densely populated with amacrine cells (INL, Pax6^+^, AC, arrowheads) and RGCs (GCL, Pax6^+^, RGC, arrowheads) across the four designated dorsoventral quadrants. In contrast, the dorsoventral gradient of reduced INL and GCL thickness in *Lhx2-Cre:Dyrk1a^f/f^*(M – P) animals led to a reduction in amacrine and RGC number (Pax6^+^, RGC/AC, arrowheads) which was most prominent in the D1 quadrant (M). (Q – X) Horizontal cells (Calbindin^+^, HC, arrowheads) in the INL of *Lhx2-Cre* (Q – T) exhibited comparable numbers across the retina. In contrast, the dorsoventral gradient of reduced INL thickness in *Lhx2-Cre:Dyrk1a^f/f^*(U – X) animals led to a reduction in horizontal cell number (Calbindin^+^, HC, arrowheads) which was most prominent in the D1 quadrant (U). Scale bar: (A – X) 25 μm. Abbreviations: AC, amacrine cell; BC, bipolar cell; D, dorsal; GCL, ganglion cell layer; HC, horizontal cell, INL, inner nuclear layer; ONL, outer nuclear layer; RGC, retinal ganglion cell; V, ventral.

**Figure 6.**
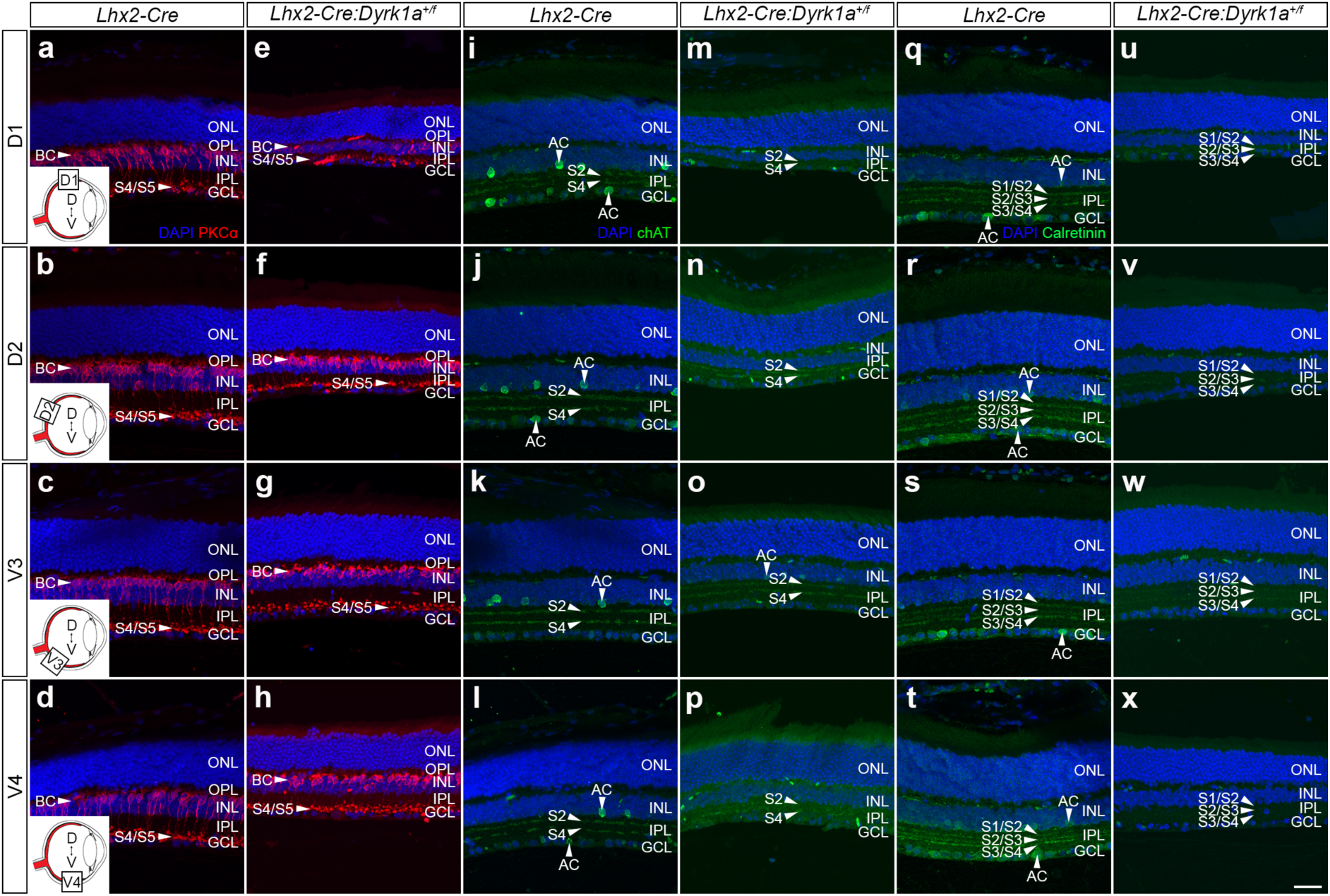
*Dyrk1a* haploinsufficiency induces inner retina stratification deficits. (A – X) Representative immunohistochemical staining of coronal eye sections taken from *Lhx2-Cre* and *Lhx2-Cre:Dyrk1a^+/f^* adult mice (6 – 8 weeks) across four designated dorsoventral quadrants (D1, D2, V3 and V4) spanning the entire retina relative to the optic nerve head. (A – H) Rod ON bipolar cell bodies (PKCα^+^, BC, arrowheads) in the INL of *Lhx2-Cre* animals (A – D) show consistent dorsoventral IPL neurite stratification patterns (S4/S5). Contrastingly, the reduced rod ON bipolar cell numbers (PKCα^+^, BC, arrowheads) in heterozygous mice (E – H) lead to irregular S4/S5 sublamina termination patterns most evident in the D1 quadrant (E) (I – P) Cholinergic amacrine cell bodies (chAT^+^, AC, arrowheads) in the INL and GCL of transgenic controls (I – L) exhibit typical neurite stratification patterns (S2/S4). In contrast, the reduced numbers of amacrine cells (chAT^+^, AC, arrowheads) in *Lhx2-Cre:Dyrk1a^f/f^*mice (M – P) lead to irregular IPL neurite stratification particularly in the D1 quadrant where S2/S4 sublamina are absent. (Q – X) Calretinin^+^ amacrine cells (AC, arrowheads) in the INL and GCL of *Lhx2-Cre* animals (Q – T) display consistent dorsoventral IPL neurite stratification patterns (S1/S2, S2/S3 and S3/S4). Contrastingly, heterozygous animals (U – X) showed diminished Calretinin^+^ amacrine cell numbers and consequent abnormal IPL neurite stratification patterns (S1/S2, S2/S3 and S3/S4) across all quadrants. Scale bar: (A – X) 25 μm. Abbreviations: AC, amacrine cell; BC, rod bipolar cell; D, dorsal; GCL, ganglion cell layer; INL, inner nuclear layer; IPL, inner plexiform layer; ONL, outer nuclear layer; OPL, outer plexiform layer; S, sublamina; V, ventral.

### Dyrk1a haploinsufficiency impairs the organisation of bipolar cell mosaics

The vertical circuit of the retina is organised along the photoreceptor-bipolar-RGC axis and facilitates the transmission of visual information from the outer to the inner retina. A ciritcal aspect of this circuit is the precise arrangement of bipolar cell mosaics which are essential for accurate spatial and contrast respresentation (Ghosh et al., 2004). Flat mounted retina were therefore analysed from transgenic control and homozygote mice (6 – 8 weeks) to determine whether the reductions in bipolar cell number disrupted their mosaic arrangement (Figure 7). Bipolar cell bodies displayed a consistent spatial distribution in *Lhx2-Cre* mice (Chx10^+^, Figure 7A – 7D). In contrast, the dorsal-most quadrant of heterozygous retinas exhibited irregular bipolar cell body positioning (Chx10^+^, Figure 7E) while the remaining regions were comparable to transgenic controls (Chx10^+^, Figure 7F – 7H). The apparent disruption of bipolar cell mosaic regularity in the dorsal-most retina of *Lhx2-Cre:Dyrk1a^+/f^* animals was confirmed by Nearest Neighbour and Voronoi domain analysis (Reese and Keeley, 2015). All dorsoventral quadrants in transgenic controls exhibited comparable spatial properties with consistent nearest neighbour (NNRI) and Voronoi domain (VDRI) regularity indices along with low coefficient of variation (CV) (Figure 7I – 7L). As expected, the dorsal-most region of heterozygous mice displayed irregular mosaic spacing as evidenced by skewed NNRI and VDRI values in addition to increased CV (Figure 7M) whereas the remaining quadrants showed spatial properties and variances comparable to transgenic controls (Figure 7N – 7P). These dorsal-most differences between *Lhx2-Cre* and *Lhx2-Cre:Dyrk1a^+/f^* animals were subsequently confirmed by quantitative (Figure 7Q – 7X) and frequency distribution analyses (Figure S4). Taken together, *Dyrk1a* haploinsufficiency selectively disrupted bipolar cell spatial organisation in a region-specific manner with the most pronounced deficits observed in the dorsal-most retina of heterozygous mice.

**Figure 7.**
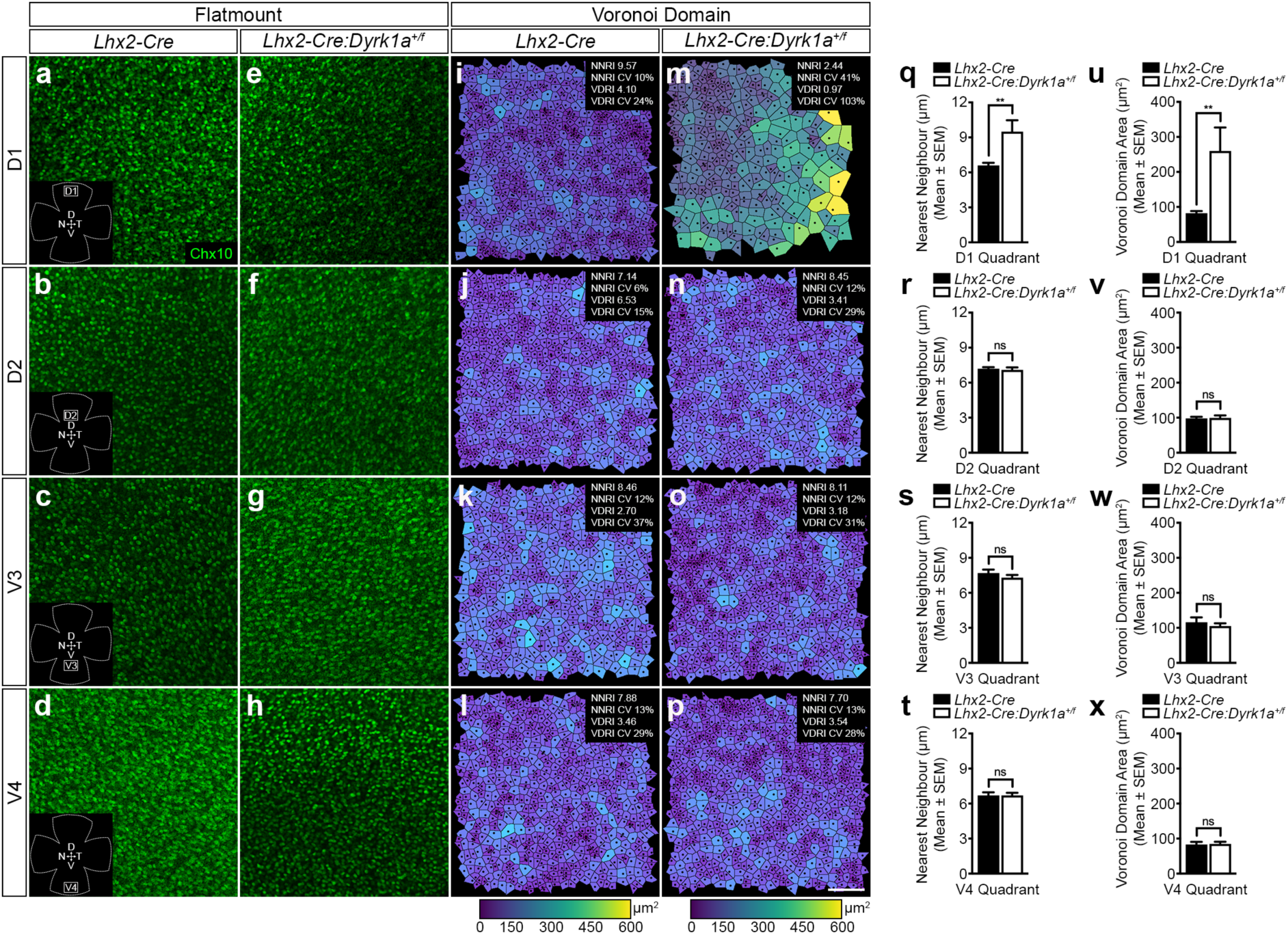
*Dyrk1a* haploinsufficiency impairs the organisation of bipolar cells mosaics. (A – H) Representative flat mount retina images from *Lhx2-Cre* (A – D) and *Lhx2-Cre:Dyrk1a^+/f^*(E – H) adult mice (6 – 8 weeks) showing the mosaic distribution of bipolar cells across four defined dorsoventral quadrants (D1, D2, V3, and V4) spanning the entire central retina relative to the optic nerve head. Transgenic controls displayed an even distribution of bipolar cells across all quadrants. In contrast, heterozygous animals showed a reduced number of Chx10^+^ bipolar cells in the dorsal-most quadrant (E) while the remaining domains (F – H) were comparable to transgenic controls. (I – P) Representative Voronoi domain plots of bipolar cell distribution in *Lhx2-Cre* (I – L) and *Lhx2-Cre:Dyrk1a^+/f^*(M – P) adult mice (6 – 8 weeks). Transgenic controls showed evenly spaced bipolar cell mosaics with nearest neighbour (NNRI) and Voronoi domain regularity indices (VDRI) indicating robust spatial regularity. In contrast, the dorsal-most quadrant of heterozygous mice exhibited mosaic spacing variability reflected by irregular Voronoi domain areas and reduced regularity indices (M) while the remaining quadrants (N – P) retained regularity indices comparable to transgenic controls. (Q – T) Quantitative analysis of nearest neighbour distance in *Lhx2-Cre* (*n* = 5) and *Lhx2-Cre:Dyrk1a^+/f^*(*n* = 8) adult animals (6 – 8 weeks). Nearest neighbour distance was significantly increased in the dorsal-most quadrant of heterozygous animals owing to a reduced cell number (Q) while the other regions remained comparable to transgenic controls (R – T). (U – X) Quantitative analysis of Voronoi domain area in *Lhx2-Cre* (*n* = 5) and *Lhx2-Cre:Dyrk1a^+/f^* (*n* = 8) adult animals (6 – 8 weeks). A significant increase in Voronoi domain area was observed in the dorsal-most quadrant of heterozygous mice owing to decreased cell number (U) while the other quadrants showed no significant differences (V – X). All data represents the mean ± SEM. Statistical differences were calculated using Mann-Whitney tests. p-values are denoted as follows: **p≤ 0.01. Scale bar: (A – P) 50 μm. Abbreviations: CV, coefficient of variation; D, dorsal; N, nasal; NNRI, nearest neighbour regularity index; T, temporal; V, ventral; VDRI, Voronoi domain regularity index.

### *Dyrk1a* haploinsufficiency leads to reduced bipolar cell activity

Electroretinography (ERG) was performed to assess whether the observed region-specific reduction in bipolar cell number and spatial organisation resulted in functional activity deficits (Figure 8). ERG waveform components consist of an initial negative deflection due to photoreceptor hyperpolarisation in response to light stimulus (a-wave) followed by a positive deflection which primarily reflects ON bipolar cell activity (b-wave) (Figure 8A). Dark-adapted scotopic ERG waveforms from transgenic control and heterozygous mice (6 – 8 weeks) displayed comparable a-wave amplitudes and latencies towards a series of increasing flash intensities that was indicative of intact photoreceptor activity (Figures 8B – 8D). In contrast, scotopic b-wave bipolar cell amplitudes were significantly reduced in *Lhx2-Cre:Dyrk1a^+/f^* animals at higher stimulus intensities (Figure 8E) albeit with similar latency to transgenic controls (Figures 8F). This also held true for light-adapted photopic ERG recordings using both UV and green flash stimuli (Figure 8G – 8J). Although a-wave amplitudes were comparable between genotypes (Figure 8G and 8I) the heterozygous mice exhibited significantly reduced photopic b-wave amplitudes at higher stimulus intensities (Figure 8H and Figure 8J). These findings indicate that photoreceptor function remains intact in *Lhx2-Cre:Dyrk1a^+/f^*mice whereas bipolar cell-driven responses across both scotopic and photopic pathways are compromised.

**Figure 8.**
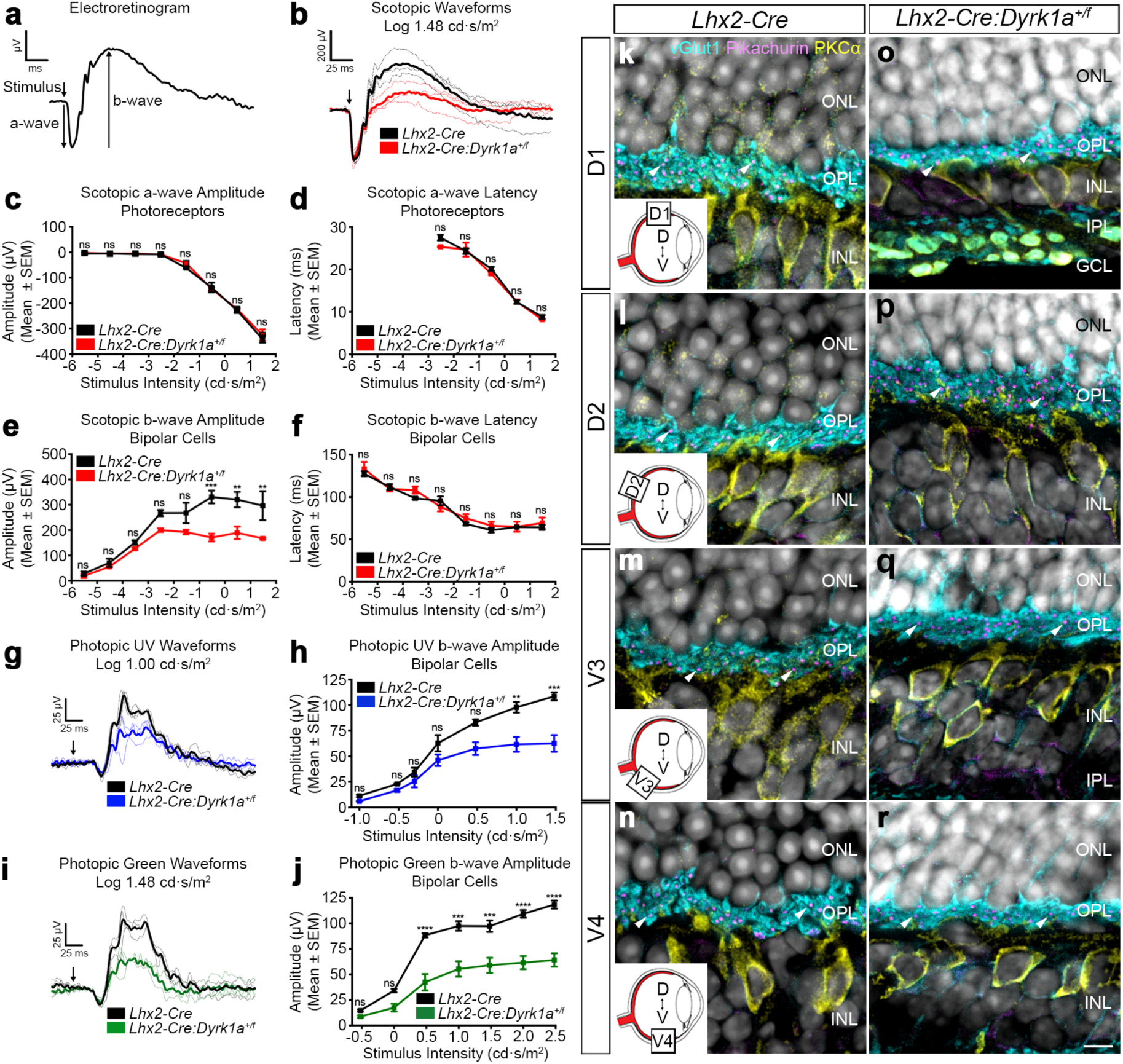
*Dyrk1a* haploinsufficiency leads to reduced bipolar cell activity. (A) Schematic representation of electroretinogram (ERG) components. An initial negative deflection (a-wave) is observed upon photoreceptor stimulation (arrow) that is followed by a subsequent positive deflection (b-wave) that corresponds to bipolar cell activity. ERG waveform quantification involves measuring the amplitude (μV) and latency (ms) of the a- and b-waves to increasing stimulus intensities (cd.s/m^2^) and thus provides an evaluation of retinal function. (B) Representative dark-adapted scotopic ERG waveforms recorded from *Lhx2-Cre* (*n* = 3) and *Lhx2-Cre:Dyrk1a^+/f^*(*n* = 3) animals (6 – 8 weeks) in response to a single flash stimulus (log 1.48 cd.s/m^2^). Comparable a-wave responses are observed for both groups while b-wave amplitudes are reduced in heterozygous animals. Thick lines represent genotype-mean responses while thin lines represent individual mice. (C – D) Quantifcation of photoreceptor amplitude (C) and latency (D) responses in *Lhx2-Cre* (*n* = 3) and *Lhx2-Cre:Dyrk1a^+/f^*(*n* = 3) animals (6 – 8 weeks) in repsonse to a series of increasing scotopic flash stimuli (log cd.s/m^2^). No significant differences are observed between the groups. (E – F) Quantification of bipolar cell amplitude (E) and latency (F) responses in *Lhx2-Cre* (*n* = 3) and *Lhx2-Cre:Dyrk1a^+/f^* (*n* = 3) animal (6 – 8 weeks) towards increasing scotopic flash stimuli (log cd.s/m^2^). Heterozygous animals have a significant reduction in bipolar cell amplitude response at higher stimulus intensities (E) albeit with similar latency (F). (G) Representative light-adapted photopic ERG waveforms recorded from *Lhx2-Cre* (*n* = 3) and *Lhx2-Cre:Dyrk1a^+/f^* (*n* = 3) animals (6 – 8 weeks) in response to a single UV flash stimulus (log 1.00 cd.s/m^2^). Comparable a-wave responses are observed for both groups while b-wave amplitudes are reduced in heterozygous animals. Thick lines represent genotype-mean responses while thin lines refer to individual mice. (H) Quantification of bipolar cell amplitude response in *Lhx2-Cre* (*n* = 3) and *Lhx2-Cre:Dyrk1a^+/f^* (*n* = 3) animal (6 – 8 weeks) in response to an increasing photopic UV stimulus intensity series (log cd.s/m^2^). Heterozygous animals have a significant reduction in bipolar cell amplitude response at higher stimulus intensities. (I) Representative light-adapted photopic ERG waveforms recorded from *Lhx2-Cre* (*n* = 3) and *Lhx2-Cre:Dyrk1a^+/f^*(*n* = 3) animals (6 – 8 weeks) in response to a single green flash stimulus (log 1.48 cd.s/m^2^). Comparable a-wave responses are observed for both groups while b-wave amplitudes are reduced in heterozygous animals. Thick lines represent genotype-mean responses while thin lines refer to individual mice. (J) Quantification of bipolar cell amplitude response in *Lhx2-Cre* (*n* = 3) and *Lhx2-Cre:Dyrk1a^+/f^*(*n* = 3) animal (6 – 8 weeks) in response to a photopic green stimulus intensity series (log cd.s/m^2^). Heterozygous animals have a significant reduction in bipolar cell amplitude response at higher stimulus intensities. (K – R) Representative immunohistochemical staining of coronal eye sections taken from *Lhx2-Cre* and *Lhx2-Cre:Dyrk1a^+/f^*adult mice (6 – 8 weeks) across four designated dorsoventral quadrants (D1, D2, V3 and V4) spanning the entire retina relative to the optic nerve head. OPL ribbon synapse assemblies appear comparable in both groups (K – R, arrowheads) and were composed of dense photoreceptor terminals (vGlut1^+^) that overlapped with well organised bipolar cell dendritic processes (PKCα^+^). Additionally, the linear alignment of all ribbon junctions (Pikachurin^+^) indicated intact photoreceptor and bipolar cell synapses. All data represents the mean ± SEM. Statistical differences were calculated using a two-way ANOVA followed by a Sidak’s multiple comparison test. p-values are denoted as follows: **p≤ 0.01, ***p≤ 0.001 and ****p≤ 0.0001. Scale bar: (K – R) 5 µm. Abbreviations: D, dorsal; GCL, ganglion cell layer; INL, inner nuclear layer; IPL, inner plexiform layer; ONL, outer nuclear layer; OPL, outer plexiform layer; V, ventral.

Immunohistochemistry to examine OPL morphology was performed to exclude whether these attenuated bipolar cell-driven response were due to aberrant photoreceptor–bipolar cell contacts. Ribbon synapse assemblies appeared comparable in both groups (Figure 8K – 8R, arrowheads) and were composed of dense photoreceptor terminals (vGlut1^+^) that overlapped with well organised bipolar cell dendritic processes (PKCα^+^). Additionally, the alignment of all ribbon junctions was maintained (Pikachurin^+^) thus indicating intact and aligned photoreceptor and bipolar cell synapses. Taken together, these observations suggest that the structural integrity of the OPL remains intact in heterozygous animals and that the attenuated bipolar cell-driven response is due to the reduced number and spatial organisation of these neurons particularly in the dorsal-most retina.

## Discussion

Using a bioinformatics approach *Dyrk1a* was identified as a novel downstream mediator of *Lhx2* transcriptional programs during mouse retinal development. Conditional deletion of *Dyrk1a* resulted in increased apoptosis and a dorsoventral gradient of reduced inner retinal thickness predominantly affecting the dorsal-most region of heterozygous animals. Specifically, *Dyrk1a* haploinsufficiency reduced bipolar cell numbers and mosaic organisation that resulted in reduced bipolar cell-driven activity (Masland, 2012). These findings therefore provide strong evidence that *Dyrk1a* acts as a transcriptional target of *Lhx2* and is a key regulator of inner retina development and circuit formation in a dosage-dependant manner.

Bipolar cell mosaics in transgenic controls exhibited nearest neighbour distances and Voronoi domain areas that varied slightly across dorsoventral quadrants. These observations indicated that bipolar cell positioning is governed by density-dependent organisation and local packing constraints (Keeley et al., 2017; Wassle et al., 2009) in contrast to the fixed exclusion model described for horizontal and amacrine cell mosaics (Kay et al., 2012; Reese et al., 2005; Wassle and Riemann, 1978). The presented data therefore support a model in which bipolar cell mosaics are primarily determined by soma size and geometric packing rather than by classical exclusion mechanisms involving homotypic repulsion and molecularly defined exclusion zones (Reese and Keeley, 2015). Moreover, the ERG recordings in *Dyrk1a* haploinsufficient mice strongly support the concept that local changes in cell number not only alter mosaic organisation but also impact bipolar cell-driven activity. Together, these findings indicate that bipolar cell mosaics are shaped predominantly by density-dependent packing constraints with local variations in cell density directly influencing both spatial arrangement and functional output.

*Dyrk1a* haploinsufficient mice exhibited marked disruption of IPL stratification including the loss of rod bipolar sublamina consistent with reduced bipolar-driven responses and ON-pathway output (Euler et al., 2014; Haverkamp and Wassle, 2000). The absence of characteristic amacrine cell termination patterns further indicated defective laminar targeting within the IPL and disorganisation of processes that normally support direction-selective circuitry (Euler et al., 2002; Famiglietti, 1983a; Famiglietti, 1983b; Haverkamp and Wassle, 2000). Since IPL lamination is essential for pathway-specific connectivity among inner retinal neurons these findings suggest that altered bipolar mosaics are accompanied by secondary defects in amacrine organization and synaptic partner integration (Masland, 2012; Sanes and Masland, 2015). These defects are primarily caused by apoptosis resulting from *Dyrk1a* haploinsufficiency which disrupts the developmental timing of bipolar cell maturation and subsequent assembly of IPL circuits. Functionally such stratification errors would weaken ON-pathway activity and degrade contrast processing and receptive field structure consistent with the reduced ERG b-wave amplitudes observed (Robson and Frishman, 2014). Thus, reduced bipolar cell number and compromised mosaic integrity provide the primary explanation for diminished retinal output in *Dyrk1a* haploinsufficient mice while the accompanying IPL disorganisation likely reflects downstream circuit assembly defects that further limit the precision of retinal signal integration and visual function (Euler et al., 2014; Masland, 2012).

A lack of rhodopsin^+^ phagosomes was observed in the dorsal retina of *Dyrk1a* haploinsufficient mice indicative of impaired rod outer segment shedding (Kevany and Palczewski, 2010; Nandrot et al., 2012). Since the *Lhx2-Cre* transgene drives target allele recombination in RPCs prior to their segregation into retina and pigment epithelium lineages (Hagglund et al., 2011) the loss of *Dyrk1a* in both cell types likely disrupted bidirectional signaling that is required for outer segment tip recognition and engulfment. The preserved rod phototransduction and normal outer segment morphology observed in heterozygous animals argues against primary defects in disc morphogenesis or phototransduction and instead supports a failure in activity-dependent renewal signaling from rods to the pigment epithelium. Given that young adult mice were employed in this study this phenotype likely represents an early event in outer retinal degeneration in which structural and functional integrity of the ONL is preserved while outer segment turnover is compromised. This phagocytic deficit is also mechanistically separable from the bipolar transmission abnormality and confirms that photoreceptor–pigment epithelium signaling is critical to ONL homeostasis (Kevany and Palczewski, 2010; Nandrot et al., 2012).

In conclusion, *Dyrk1a* is identified a novel downstream effector of *Lhx2*-regulated transcriptional hierarchies and this molecular axis is essential for inner retinal development and function. Notably, *Dyrk1a* haploinsufficiency increased apoptosis resulting in a dorsoventral loss of bipolar cell numbers and inner retina activity. Taken together these results establish *Dyrk1a* as a critical regulator of retinogenesis with precise gene-dosage required for visual system development and function (Cai et al., 2023; Mejecase et al., 2021). These findings also have broader implications for understanding how transcriptional hierarchies and gene-dose-dependent regulation orchestrate CNS development and connectivity.

## Materials and Methods

### Bioinformatics

Bioinformatic analysis was conducted in the R environment (version 4.3.3) using previously published NCBI GEO datasets (GSE172457 and GSE75889) (de Melo et al., 2016; Li et al., 2022). Data normalization and differential expression analysis were conducted using DESeq2 (version 1.42.1) (Love et al., 2014) (p<0.05 and a minimum row sum of >10 reads). Intersection and gene ontology (GO) analyses of differentially expressed genes (DEGs) (padj <0.05, baseMean >10, log2FC <-0.3) was performed in ClusterProfiler (version 4.10.1) (Wu et al., 2021). Gene enrichment analyses were performed using EnrichR (version 3.2) (Kuleshov et al., 2016). Analyses were aligned to the GRCm38 mouse genome.

### Animals

Animal experiments were approved by the Animal Review Board at the Court of Appeal of Northern Norrland (#A22-2023) and the Finnish Project Authorisation Board (#ESAVI/37754/2024). The derivation and genotyping of *Tg(Lhx2-Cre)1Lcar* (referred to as *Lhx2-Cre*), *Dyrk1a^tm1Jdc/J^* (referred to as *Dyrk1a^+/f^* or *Dyrk1a^f/f^*) and *Gt(ROSA)26Sor^tm1sor^*(referred to as *ROSA26R^R/R^*) mice have been previously described (Hagglund et al., 2011; Soriano, 1999; Thompson et al., 2015). Genotypes were determined by PCR analysis of genomic DNA extracted from ear biopsies (Sambrook and Russell, 2001). The morning of the mating plug was considered as E0.5. Experimental analyses were conducted using both males and females on a *129/Sv:CBA:C57BL/6:NMRI:DBA2/J* mixed genetic background. Littermates carrying the *Dyrk1a^tm1Jdc/J^* allele were used as controls for lineage tracing analysis while littermates that carried the *Lhx2-Cre* transgene were used as controls for all other experiments. Mice were sedated by an intraperitoneal injection of either (i) medetomidine (0.75 mg/kg), midazolam (5 mg/kg) and fentanyl (0.05 mg/kg) prepared in saline or (ii; for *in vivo* experiments) ketamine (60 mg/kg) and medetomidine (0.4 mg/kg) prepared in saline. Anaesthesia was reversed by the administration of an antidote consisting of atipamezole (2.5 mg/kg), flumazenil (0.5 mg/kg) and naloxone (1.2 mg/kg) prepared in saline. During sedation and recovery all mice were kept in their home cages on heating pads.

### In Situ Hybridisation

A heated needle was first used to burn an orientation mark at the dorsal pole of the eye prior to enucleation in all postnatal animals. Embryonic heads or enucleated eyes were fixed in 4% (w/v) PFA in PBS for 2 h on ice and equilibrated in 30% (w/v) sucrose in PBS overnight at 4°C. Tissues were embedded in dorsoventral orientations in OCT compound (Sakura Finetek). *In situ* hybridisation on cryosections (10 μm) was performed as previously described (Schaeren-Wiemers and Gerfin-Moser, 1993). The complete open reading frame of *Dyrk1a* (ENSMUST00000119878, nucleotide position 1 – 2292 bp) was used to generate a riboprobe.

### Immunohistochemistry

A heated needle was first used to burn an orientation mark at the dorsal pole of the eye prior to enucleation in all postnatal animals. For cryo- or paraffin section analyses the embryonic heads or enucleated eyes were first fixed in 4% (w/v) PFA in PBS for 2 h on ice. For cryosection preparation the fixed tissues were equilibrated overnight in 30% (w/v) sucrose in PBS at 4°C and then embedded in dorsoventral orientation in OCT (Sakura Finetek). For paraffin sectioning the fixed tissues were immersed in 70% (v/v) ethanol at 4°C overnight and then paraffin embedded in a dorsoventral orientation. For whole mount analyses a heated needle was first used to burn an orientation mark at the dorsal pole of the eye prior to enucleation. The eye was then placed in a 48-well plate and fixed in 4% (w/v) PFA in PBS for 10 min on ice. The cornea and lens were then removed and an incision was made at the dorsal pole of the eye cup to maintain orientation. The eye cup was then fixed for a further 30 min in 4% (w/v) PFA in PBS on ice. The retina was subsequently peeled away from the pigment epithelium and four incomplete radial incisions were made along the dorsal, temporal, ventral and nasal axes to yield four petals attached to one another at the central optic disc region. A separate incision was also placed in the dorsonasal petal for orientation purposes. Immunohistochemistry was performed on sections (10 μm) or whole mount retina as previously described (Cantrup et al., 2012). For all experiments involving paraffin sections the samples were first cleared by a 2 x 10 min incubation in xylenes followed by rehydration through a graded series of ethanol (99.5%, 95%, 70% and 50% (v/v) in PBS) and antigen retrieval (Fitter et al., 2017). An additional blocking step involving M.O.M. Blocking Reagent (Vector Labs) was used in experiments involving monoclonal primary antibodies. The following primary antibodies and dilutions were used in this study: Calbindin (1:300, Sigma-Aldrich, #C9848); Calretinin (1:1000, Swant, #CG1); cleaved Caspase3 (Asp175) (1:500, Cell Signalling, #9661S); chAT (1:100, Millipore, #AB144P); Chx10 (1:50, SCBT, #SC365519); Dyrk1a (1:100, Novus, #NBP1-84032); Isl1/2 (recognises both Isl1 and Isl2 proteins) (1:500; DSHB, #39.4D5); M-opsin (1:500, Proteintech, #30975-1-AP); Pax6 (1:100, DSHB, #Pax6); Pikachurin (1:200, Proteintech, #14578-1-AP); Rhodopsin (1:500, Millipore, #MABN15); S-opsin (1:500, Proteintech, #24660-1-AP); TH (1:300, Millipore, #AB152); vGlut1 (1:2000, Millipore, #AB5905).

### Lineage Tracing Analysis

A heated needle was first used to burn an orientation mark at the dorsal pole of the eye prior to enucleation and the eyes were fixed in 4% (w/v) PFA in PBS for 30 min on ice. The tissues were then equilibrated overnight at 4°C in 30% (w/v) sucrose in PBS and embedded in a dorsoventral orientation in OCT compound (Sakura Finetek). Lineage tracing analyses were performed on cryosections (10 μm) as described previously (Hagglund et al., 2011).

### Quantitative Polymerase Chain Reaction (qPCR)

Enucleated eyes from postnatal animals were dissected and the retina was snap frozen in LN2. Genomic DNA was extracted from individual retina (Sambrook and Russell, 2001) and used as the template for qPCR using the SsoAdvanced™ Universal SYBR® Green Supermix (Bio-Rad) and CFX Connect™ Real-Time System (Bio-Rad). Amplification reactions were performed in triplicate and contained genomic DNA template (20 ng) and oligonucleotide pairs that amplified across exon 4 (R4F 5′-CTGTGGTAGTGGGTGTTCAATA-3′ and R4R 5′-CCGCCTCTGTAACATGACAA-3′) and exon 6 (T6F 5′-GGAGGAGGATGTAAACAGTGAG-3′ and T6R 5′-TCTTGCTCCACTCTGTCATAAG-3′) of the mouse *Dyrk1a* gene (ENSMUSG00000022897). For quantification analysis the level of product generated by the *LoxP* target primer pair (exon 6) was normalised against the level of product amplified by the reference primer pair (exon 4) for each biological replicate.

### Immunoblotting

Enucleated eyes from postnatal animals were dissected to remove the lens and cornea before snap freezing the eye cup in LN2. Tissues were mechanically homogenised in RIPA Extraction Buffer (Thermo Fisher Scientific) containing protease and phosphatase inhibitors (Complete Mini & PhosSTOP, Roche). The concentration of the soluble protein fraction was measured using the BCA Protein Assay Kit (Thermo Fisher Scientific). Immunoblotting was performed on protein extracts (30 µg) using Criterion TGX gels and nitrocellulose membranes (Bio-Rad). Reactive proteins were visualised using SuperSignal West Dura Extended Duration Substrate (Thermo Fisher Scientific). Image capture was performed using a ChemiDoc MP Imaging System (Bio-Rad). The following primary antibodies and dilutions were used: β-actin (1:20000, Sigma Aldrich, #A3854); cleaved Caspase3 (Asp175) (1:1000, Cell Signalling, #9661S); Dyrk1a (1:1000, Abnova, #H00001859). For quantification analysis the intensity of the Dyrk1a (90 kDa) and cleaved Caspase3 (19/17 kDa) bands were normalised against the intensity of the β-actin (42 kDa) band for each biological replicate.

### Optical Coherence Tomography (OCT)

OCT imaging was conducted using a Micron IV Retina Imaging System (Phoenix Research Laboratories). Mice were anesthetised and Carbomer eye gel (2 mg/g) was then applied onto each cornea. A stage holding the animal was stereoscopically adjusted to centre the field of view at the optic nerve head. B-scans of the retina were subsequently captured at cardinal (dorsal, ventral, nasal, temporal) and intercardinal (dorsonasal, dorsotemporal, ventronasal, ventrotemporal) orientations. Retinal layer thickness analyses were performed at fixed distance (500 µm) from the optic nerve head.

### Histology

For histology analyses a heated needle was first used to burn an orientation mark at the dorsal pole of the eye prior to enucleation. Enucleated eyes were fixed in 4% (w/v) PFA in PBS for 2 h on ice and then postfixed in 2% (v/v) glutaraldehyde in PBS overnight at 4°C. Eyes were then immersed in 70% (v/v) ethanol at 4°C overnight and paraffin embedded in a dorsoventral orientation. Paraffin sections (10 μm) were cleared by 2 x 10 min incubation in xylenes before rehydration though a graded series of ethanol (99.5%, 95%, 90% and 80% (v/v) in PBS). Haematoxylin and eosin staining was performed as previously described (Hagglund et al., 2011).

### Electroretinogram (ERG)

Animals were dark-adapted overnight and all handling during scotopic ERG recordings were performed under red light conditions. Pupils were first dilated by the topical addition of 10% (w/v) phenylephrine and 5% (v/v) tropicamide. The mice were subsequently anesthetised and placed on a heated stage. Carbomer eye gel (2 mg/g) was then applied to the corneas and rod-driven scotopic ERG recordings (Pazur et al., 2024) were performed using an Espion E3 ERG device (Diagnosys LLC) using an in-house scripted protocol (Diagnosys Espion V6 Software) that contained 8 different monochromatic green light stimulation steps: Step 1, stimulus intensity 0.000003 cd·s/m^2^, 20 sweeps, 2 s interstimulus interval (ISI) between sweeps; Step 2, 0.00003 cd·s/m^2^, 15 sweeps, 3 s ISI; Step 3: 0.0003 cd·s/m^2^, 15 sweeps, 3 s ISI; Step 4: 0.003 cd·s/m^2^, 12 sweeps, 8 s ISI; Step 5: 0.03 cd·s/m^2^, 6 sweeps, 15 s ISI; Step 6: 0.3 cd·s/m^2^, 4 sweeps, 15 s ISI; Step 7: 3 cd·s/m^2^, 2 sweeps, 45 s ISI; and Step 8, 30 cd·s/m^2^, 2 sweeps, 60 s ISI. Following scotopic ERG recording the animals were moved into standard laboratory light conditions. To produce cone-driven photopic ERGs the light flashes were superimposed to a steady background light at 20 cd/m^2^ using a Celeris Rodent ERG System (Diagnosys LLC). UV (peak emission 370 nm, bandwidth 50 nm) and green (peak emission 544 nm, bandwidth 160 nm) monochromatic LEDs were used and stimulation was produced using an in-house scripted protocol (Diagnosys Espion V6 Software) using the following stimulation steps: Step 1, green 0.3 cd·s/m^2^, 40 sweeps, 200 ms ISI; Step 2, green 1 cd·s/m^2^, 30 sweeps, 300 ms ISI; Step 3, UV 0.1 cd·s/m^2^, 40 sweeps, 300 ms ISI; Step 4, UV 0.3 cd·s/m^2^, 30 sweeps, 300 ms ISI; Step 5, UV 0.5 cd·s/m^2^, 25 sweeps, 300 ms ISI; Step 6, green 3 cd·s/m^2^, 15 sweeps, 300 ms ISI; Step 7, green 10 cd·s/m^2^, 15 sweeps, 300 ms ISI; Step 8, UV 1 cd·s/m^2^, 15 sweeps, 1 s ISI; Step 9, UV 3 cd·s/m^2^, 15 sweeps, 1 s ISI; Step 10: green 30 cd·s/m^2^, 15 sweeps, 1 s ISI; Step 11: UV 10 cd·s/m^2^, 10 sweeps, 1.5 s ISI; Step 12: green 100 cd·s/m^2^, 10 sweeps, 1.5 s ISI; Step 13: UV 30 cd·s/m^2^, 10 sweeps, 1.5 s ISI; and Step 14, green 300 cd·s/m^2^, 10 sweeps, 1.5 s ISI. ERG signals were acquired at sampling rate of 2 kHz and filtered using low (0.25 Hz) and high (300 Hz) frequency cutoffs. A-wave amplitudes were measured from signal baseline to the negative trough while b-wave amplitudes were calculated as the difference between the a-wave trough and the peak of the b-wave. Response peak times were measured as the latency to the respective maximum trough (a-wave) or maximum peak (b-wave). Only b-waves were analysed from the photopic ERG.

### Image Analyses

Microscopic images were captured using SP8 Falcon confocal (Leica) and an Axioscan Z1 slide scanner (Zeiss). Image analyses were performed blind to genotype. Isl1/2 and cleaved Capsase3 cell counts were performed using a custom pipeline configured in CellProfiler v4.2.8 (Carpenter et al., 2006; Kamentsky et al., 2011). Histology and flat mount analysis were performed using custom scripts written in Fiji v2.16.0/1.54r (Schindelin et al., 2012). Custom pipeline and script workflows are fully described in the Supplementary Information.

## Statistical Analyses

Statistical analyses were performed blind to genotype using Prism10 (GraphPad Software). Error bars in all figures represent the standard error of the mean (SEM). The number of animals analysed (*n*) and the statistical tests employed are given in the respective figure legend. p-values are indicated as follows: *p≤0.05, **p≤0.01, ***p≤0.001, ****p≤ 0.0001.

## Data Availability Statement

Raw data and imaging scripts are available from the corresponding author upon request.

## Author Contribution Statement

Conceptualisation: IJ, LC; Methodology: CN, AT, ASM, HL, IJ; Data Analysis: CN, AT, ASM, HL, SW, LC, IJ; Writing Original Draft: IJ; Writing Review and Editing: CN, AT, ASM, HL, SW, LC, IJ; Supervision: HL, SW, LC; Project Administration: HL, SW, LC; Funding Acquisition: HL, SW, LC.

## Acknowledgements

The Pax6 monoclonal antibody was deposited to the DSHB by Atsushi Kawakami. The Isl1/2 monoclonal antibody was deposited to the DSHB by Thomas Jessell and Susan Brenner-Morton. The authors acknowledge the Biochemical Imaging Centre at Umeå University and the National Microscopy Infrastructure (NMI) (#VR-RFI 2023-00163) for providing microscopy support.

