## Supplementary Information for "*Dyrk1a* gene dosage controls bipolar cell development and retinal connectivity"

**Running Title:** ***Dyrk1a* gene dosage and retinal connectivity.**

Christoffer Nord[1], Amol Tandon[1], Anthi-Styliani Makiou[2], Henri Leinonen[2], Sara Ivy Wilson[1], Leif Carlsson[1] & Iwan Jones[*1].

[1]Department of Medical and Translational Biology, Umeå University, Umeå, Sweden.

[2]School of Pharmacy, Faculty of Health Sciences, University of Eastern Finland, Kuopio, Finland.

**Keywords:** *Dyrk1a*, retina, bipolar cells, haploinsufficiency, connectivity, nervous system.

| | |
|---|---|
| Supplementary Figure 1 | Page 2 |
| Supplementary Figure 2 | Page 4 |
| Supplementary Figure 3 | Page 5 |
| Supplementary Figure 4 | Page 6 |
| Supplementary Table 1 | Page 7 |
| Supplementary Materials and Methods | Page 8 |
| References | Page 10 |

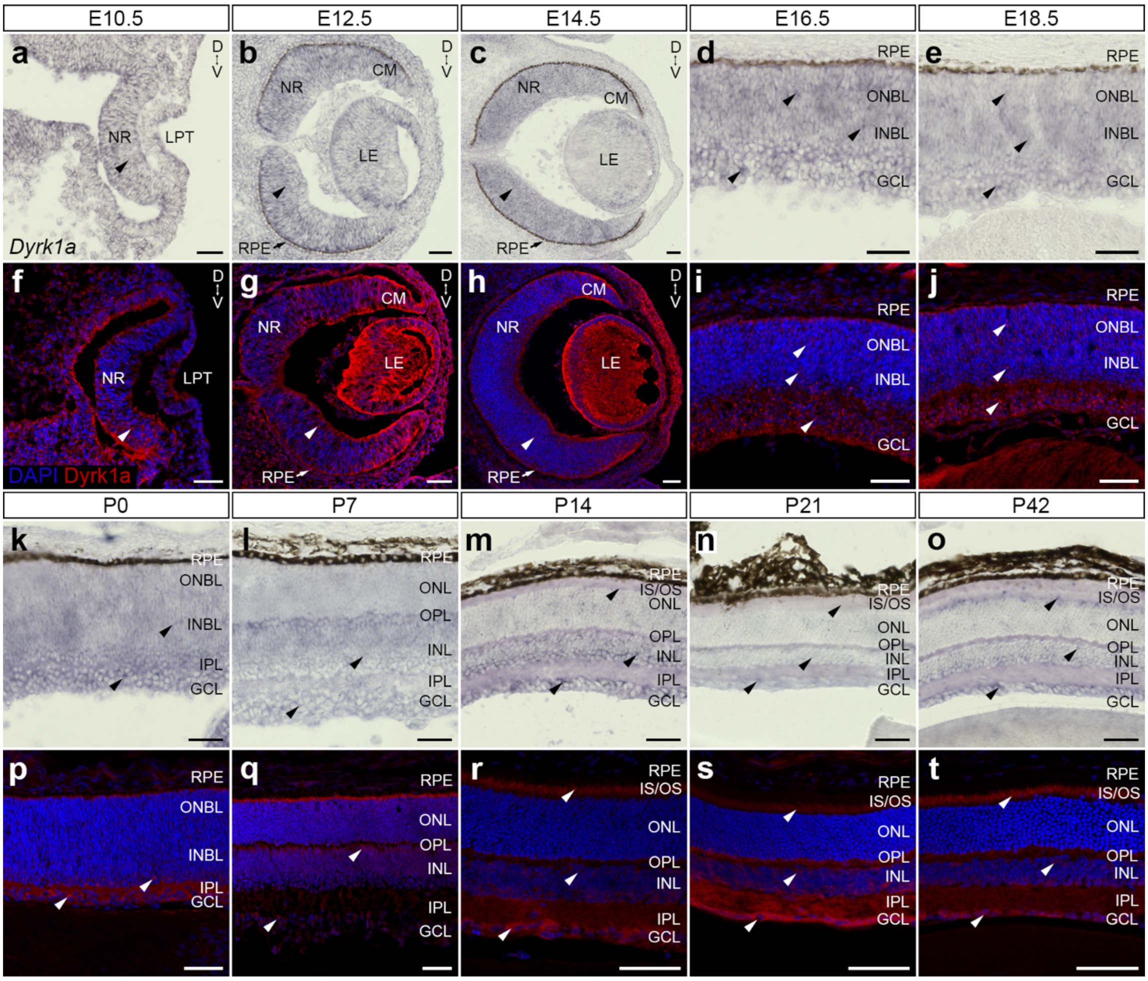


**Figure S1. Temporospatial expression of *Dyrk1a* during mouse eye development**. (A – J) Representative *in situ* hybridization (A – E) and immunohistochemistry (F – J) analyses of coronal eye sections from wild-type mice during embryonic development (E10.5 – E18.5). *Dyrk1a* transcripts were broadly distributed throughout the retina at early embryonic ages (E10.5 – E14.5, A – C, arrowheads) and were later detected in the ONBL, INBL and GCL (E16.5 – E18.5, D – E, arrowheads). Dyrk1a protein was initially observed in the ventral retina (E10.5, F, arrowhead) but subsequently became enriched throughout the retina (E12.5 – E14.5, G – H, arrowheads). At later ages Dyrk1a was uniformly distributed across the ONBL, INBL and GCL (E16.5 – E18.5, I – J, arrowheads). (K – T) Representative *in situ* hybridization (K – O) and immunohistochemistry (P – T) analyses of coronal eye sections from wild-type mice during postnatal development (P0 – P42). *Dyrk1a* transcripts were first detected in the INBL and GCL (P0, K, arrowheads) and subsequently accumulated in the INL and GCL (P7, L, arrowheads). At later ages *Dyrk1a* expression became localised to the photoreceptor segments, INL and GCL (P14 – P42, M – O, arrowheads). Protein distribution again mirrored gene expression with immunoreactivity first observed in the INBL and GCL (P0, P, arrowheads) and subsequently in the apical INL and GCL (P7, Q, arrowheads). At later ages Dyrk1a was localised to the photoreceptor segments, INL and GCL (P14 – P42, R – T, arrowheads). Scale bars: (A – T) 50 µm. Abbreviations: CM, ciliary margin; D, dorsal; GCL, ganglion cell layer; INBL, inner neuroblastic layer; INL, inner nuclear layer; IPL, inner plexiform layer; IS/OS, inner and outer segments; LE, lens; LPT, lens pit; NR, neural retina; ONBL, outer neuroblastic layer; ONL, outer nuclear layer; OPL, outer plexiform layer; RPE, retinal pigment epithelium; V, ventral.

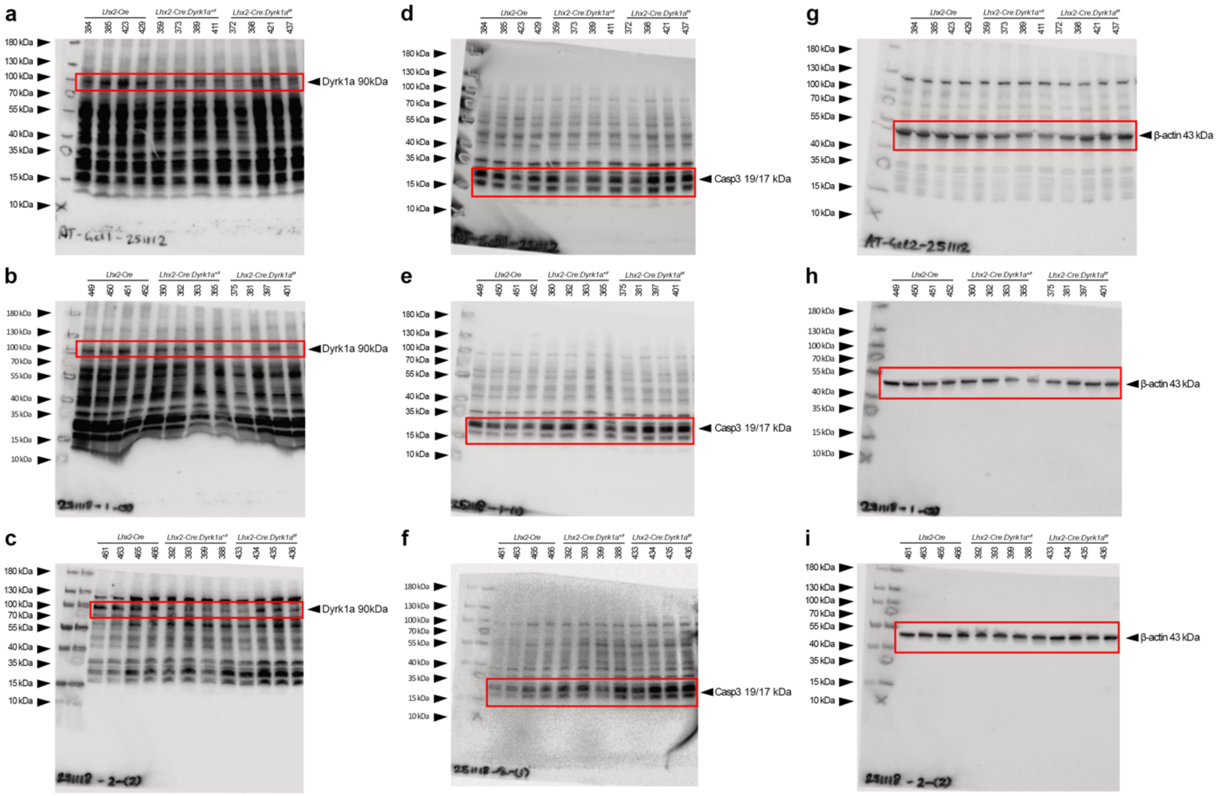


**Figure S2.** (A – I) Immunoblotting analyses showing Dyrk1a (90 kDa, A – C), cleaved Caspase3 (19/17 kDa, D – F) and β-actin (43 kDa, G – I) levels in soluble protein extracts prepared from the retina and pigment epithelium of *Lhx2-Cre* ($n$ = 12), *Lhx2-Cre:Dyrk1a*$^{+/f}$ ($n$ = 12) and *Lhx2-Cre:Dyrk1a*$^{f/f}$ ($n$ = 12) mice at P0. Each immunoblot contains four biological replicates for each genotype. The genotype and corresponding animal identification numbers are shown above each immunoblot. Red boxes demarcate the regions employed for the quantification analyses presented in Figure 2.

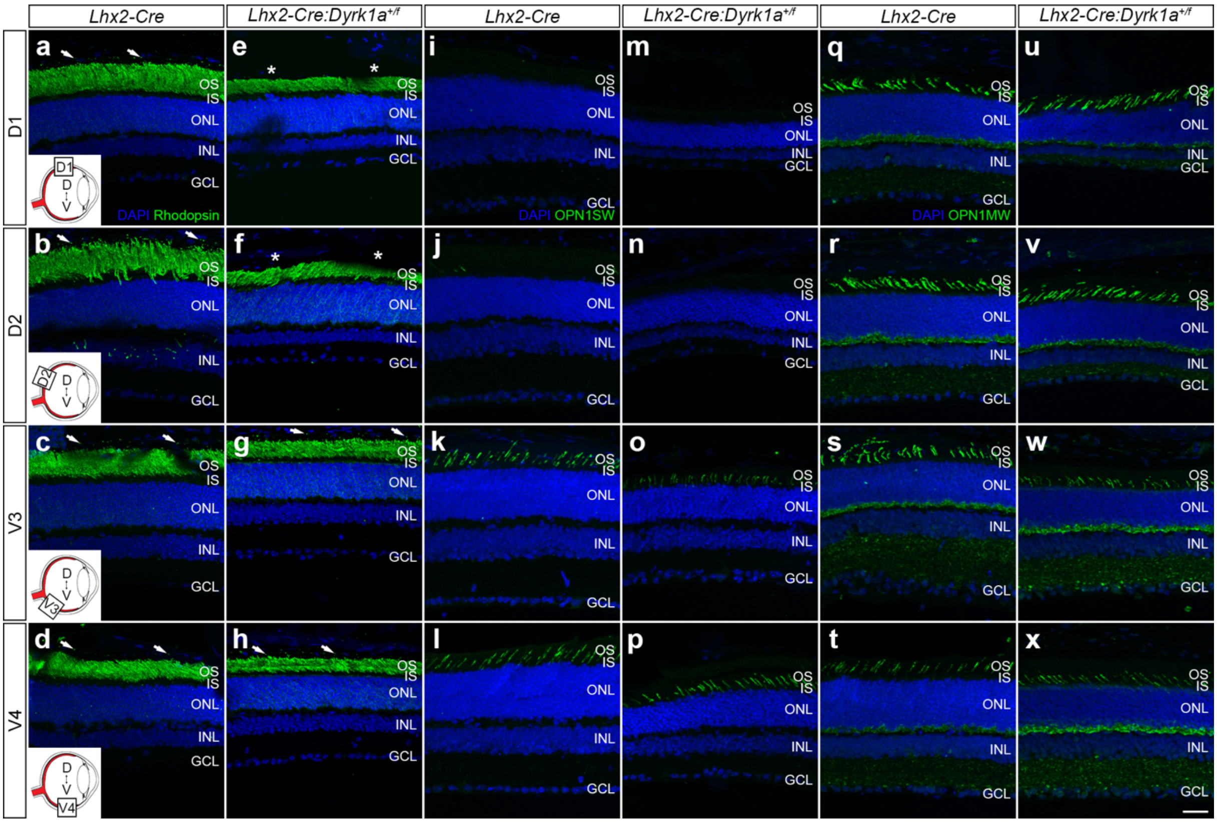


**Figure S3.** ***Dyrk1a* haploinsufficiency does not influence photoreceptor integrity.** (A – X) Representative immunohistochemical staining of coronal eye sections taken from *Lhx2-Cre* and *Lhx2-Cre:Dyrk1a*$^{+/f}$ adult mice (6 – 8 weeks) across four designated dorsoventral quadrants (D1, D2, V3 and V4) spanning the entire retina relative to the optic nerve head. The ONL of both groups exhibited densely packed photoreceptor cell bodies (DAPI$^{+}$, A – X) and photoreceptor segments enriched in rhodopsin photopigment (A – H). Discrete rhodopsin puncta were also detected within the pigment epithelium of *Lhx2-Cre* mice consistent with rod outer segment phagocytosis (A – D, arrows). In contrast, *Lhx2-Cre:Dyrk1a*$^{+/f}$ animals exhibited a marked reduction of phagosomes selectively in the dorsal retina (E – F, asterisks) while the ventral quadrants appeared comparable to controls (G – H, arrows). (I – X) Both groups possessed characteristic dorsal-low to ventral-high gradient of short-wave opsin (I – P) and dorsal-high to ventral-low distribution of medium-wave opsin (Q – X). Scale bar: (A – X) 25 µm. Abbreviations: D, dorsal; GCL, ganglion cell layer; INL, inner nuclear layer; IPL, inner plexiform layer; IS, inner segments; ONL, outer nuclear layer; OS, outer segments; OPL, outer plexiform layer; OPN1MW, medium wave opsin; OPN1SW, short wave opsin; V, ventral.

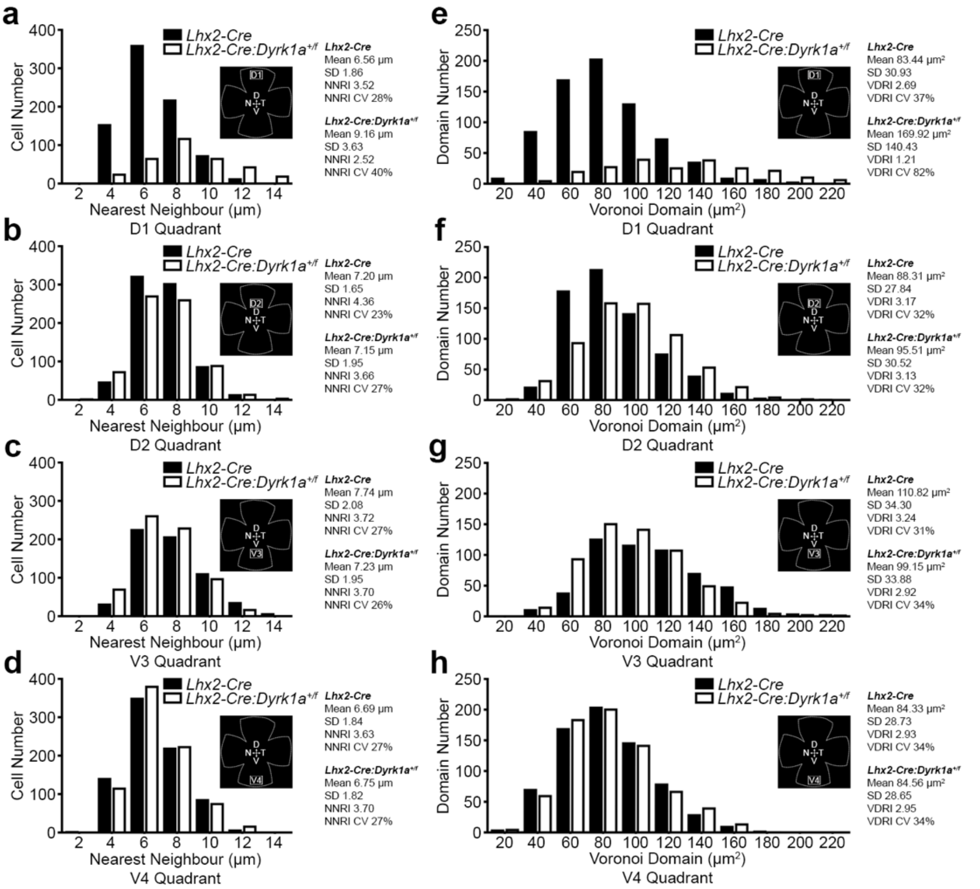


**Figure S4. *Dyrk1a* haploinsufficiency leads to irregular bipolar cell distribution.** (A – H) Retina were harvested from *Lhx2-Cre* (*n* = 5) and *Lhx2-Cre:Dyrk1a*$^{+/f}$ (*n* = 7) adult mice (6 – 8 weeks) and the spatial properties of bipolar cell mosaics across four defined dorsoventral quadrants (D1, D2, V3, and V4) spanning the entire central retina relative to the optic nerve head were determined. (A – D) Histogram frequency plots of nearest neighbour distance. Transgenic controls (A – D, solid bars) demonstrate Gaussian bipolar cell body distribution and equivalent mean nearest neighbour values across all domains. In contrast, a right-skewed bipolar cell body distribution and consequent increase in the mean nearest neighbour value was observed in the dorsal-most domain of heterozygous animals (A, open bars) while the remaining quadrants were comparable to *Lhx2-Cre* animals (B – D, open bars). (E – H) Histogram frequency plots of Voronoi domain area. Transgenic controls (A – D, solid bars) demonstrate gaussian distribution and similar mean Voronoi domain area values across all domains. In contrast, a uniform distribution curve and consequent increase in the mean Voronoi domain area was observed in the dorsal-most domain of heterozygous animals (E, open bars) while the remaining quadrants were comparable to *Lhx2-Cre* animals (F – H, open bars). Abbreviations: CV, coefficient of variation; D, dorsal; N, nasal; NNRI, nearest neighbour regularity index; T, temporal; V, ventral; VDRI, Voronoi domain regularity index.

**Table S1. Mendelian frequencies recorded during postnatal development (P0, P7, P21).**

| **Age** | ***Dyrk1a$^{+/f}$*** | ***Dyrk1a$^{f/f}$*** | ***Lhx2-Cre:Dyrk1a$^{+/f}$*** | ***Lhx2-Cre:Dyrk1a$^{f/f}$*** | **Total (*n*)** |
|---|---|---|---|---|---|
| P0 | 13 | 5 | 5 | 12 | 35 |
| P7 | 5 | 6 | 3 | 0 | 14 |
| P21 | 31 | 24 | 18 | 0 | 73 |

Mendelian frequencies were recorded from *Lhx2-Cre:Dyrk1a$^{+/f}$* (♂) and *Dyrk1a$^{f/f}$* (♀) crosses ($n = 21$).

**Supplementary Materials and Methods.**

***Cell count analysis.***

Fluorescent images of cryosections (10 μm) were captured on an SP8 Falcon confocal (Leica) and analysed using CellProfiler v4.2.8 (Carpenter et al., 2006; Kamentsky et al., 2011). An ROI mask was manually drawn on the original image using the IdentifyObjectsManually module to isolate the retina and exclude the pigment epithelium and lens. The original image was then split into individual grayscale channels (DAPI, Isl1/2 and cleaved Caspase3) and the ROI mask was applied to each channel to define the analysis area. Positively labelled nuclei in each masked channel were identified using the IdentifyPrimaryObjects module. The RelateObjects module was subsequently used to associate each Isl1/2- or cleaved Caspase3-positive object with a DAPI-positive object so that only double-positive objects were quantified. All values were exported to Prism10 for statistical analysis and the data is presented in Figure 2G – 2J.

***Histological analysis.***

Haematoxylin and eosin-stained paraffin sections (10 μm) were scanned using an Axioscan Z1 slide scanner (Zeiss). From these overview images the dorsal and ventral regions of the retina were identified according to the orientation mark burnt at the dorsal pole during dissection. A curved line following the pigment epithelium was drawn from the dorsal ciliary margin through the optic nerve head to the ventral ciliary margin. The distance of the curved line was determined and four ROIs (200 $\mu m^2$ each) were placed equidistanly along this curved line (D1, D2, V3, V4). Images were captured at each ROI and analysed in Fiji v2.16.0/1.54r (Schindelin et al., 2012). Images were segmented using Trainable Weka Segmentation to identify the nuclear layers (ONL, INL and GCL) and then converted to binary masks. Skeletonisation (2D/3D) was then applied to the binary masks to define the borders of each nuclear layer. Four vertical reference lines were then drawn equidistant across the binary mask and individual layers thickness (ONL, OPL, INL, IPL) was determined using the Fiji measure tool at each vertical line. All measurements were exported to Prism10 for statistical analysis and the data is presented in Figure 4O – 4R.

***Flat mount analysis.***

Fluorescent flat mounted retinae were scanned on an SP8 Falcon confocal (Leica). From the overview images the dorsal and ventral regions were identified according to the orientation incisions performed during dissection. A straight line was drawn from the dorsal periphery through the optic nerve head to the ventral periphery. The distance of the straight line was determined and four ROIs (300 $\mu m^2$ each) were placed equidistanly along this line (D1, D2, V3, V4). Single-plane confocal images captured at each ROI were analysed in Fiji v2.16.0/1.54r (Schindelin et al., 2012). Images were first processed using contrast-limited adaptive histogram equalization (CLAHE; block size = 127 pixels, histogram = 256 bins, maximum slope = 3) to enhance local contrast and improve visualisation of cell bodies prior to segmentation. Object segmentation was performed using Otsu automatic thresholding and thresholded images were converted to binary masks. Touching objects in the binary masks were separated using Watershed Segmentation. Seed points for each object were then identified using the Ultimate Points algorithm in combination with Euclidean distance map (EDM) to ensure a single well-defined central point-per-object. Object centroids were determined using Analyze Particles (excluding edge-touching objects) and the generated coordinates used for downstream spatial analyses. Nearest neighbour distances were calculated from the object centroid coordinates using the Nearest Neighbour Distance (NND) algorithm. Voronoi tessellations were generated from the object centroid coordinates and Voronoi Domain areas were quantified using Analyze Particles (excluding edge-touching domains). Composite spatial maps were generated by combining the Nearest Neighbour and Voronoi tesselation images using Image Calculator. All measurements were exported to Prism10 for statistical analysis and the data is presented in Figure 7I – 7X and Figure S4.

**References.**

**Carpenter, A. E., Jones, T. R., Lamprecht, M. R., Clarke, C., Kang, I. H., Friman, O., Guertin, D. A., Chang, J. H., Lindquist, R. A., Moffat, J., et al.** (2006). CellProfiler: image analysis software for identifying and quantifying cell phenotypes. *Genome biology* **7**, R100.

**Kamentsky, L., Jones, T. R., Fraser, A., Bray, M. A., Logan, D. J., Madden, K. L., Ljosa, V., Rueden, C., Eliceiri, K. W. and Carpenter, A. E.** (2011). Improved structure, function and compatibility for CellProfiler: modular high-throughput image analysis software. *Bioinformatics* **27**, 1179-1180.

**Schindelin, J., Arganda-Carreras, I., Frise, E., Kaynig, V., Longair, M., Pietzsch, T., Preibisch, S., Rueden, C., Saalfeld, S., Schmid, B., et al.** (2012). Fiji: an open-source platform for biological-image analysis. *Nature methods* **9**, 676-682.
